# Flexible use of memory by food-caching birds in a laboratory behavioral paradigm

**DOI:** 10.1101/2021.04.22.441012

**Authors:** Marissa C. Applegate, Dmitriy Aronov

**Affiliations:** Mortimer B. Zuckerman Mind Brain Behavior Institute, Columbia University, New York, NY, 10027, USA

**Author notes:** Lead contact: Dmitriy Aronov.

## Abstract

Memory is used by animals to influence navigational decisions, and neural activity in memory-related brain regions correlates to spatial variables. However, navigation is a rich behavior that contains a mix of memory-guided and memory-independent strategies. Disentangling the contribution of these strategies to navigation is therefore critical for understanding how memory influences behavioral output. To address this issue, we studied the natural spatial behavior of the chickadee, a food-caching bird. These birds hide food items at concealed, scattered locations and retrieve their caches later in time. We designed an apparatus that allows chickadees to cache and retrieve food while navigating in a laboratory arena. This apparatus enabled detailed, automated, and high-throughput tracking of key behavioral variables – including caches, retrievals, and investigations of cache sites. We built probabilistic models to fit these behavioral data using a combination of various mnemonic and non-mnemonic factors. We find that chickadees use some navigational strategies that are independent of cache memories, including opportunistic foraging and spatial biases. They combine these strategies with spatially precise and long-lasting memories of which sites contain caches and which sites they have previously checked and found to be empty. These memories are used in a context-dependent manner. During caching, chickadees avoid sites that already contain food. During retrieval, they are instead attracted to such occupied sites. These results show that a single memory can be used flexibly by a chickadee to achieve at least two unrelated behavioral goals. Our apparatus enables studying this flexibility in a tractable spatial paradigm.

## Introduction

The study of episodic memory in animals has greatly benefitted by experiments that involve spatial navigation.^1–3^ Whereas memory use is internal to the animal and not directly observable, navigational decisions that require memory are overt and trackable. In addition, memory-related neural circuits, like the hippocampus, exhibit well-described firing patterns that correlate to navigational variables – including place and head direction.^4,5^ However, navigation is a rich behavior that includes both mnemonic and non-mnemonic spatial strategies. For example, an animal using memory to obtain a reward may also engage in opportunistic foraging, use memory-independent search strategies, and exhibit spatial biases.^6–10^ It is critical to tease apart these contributions to behavior in order to eventually understand the underlying neural mechanisms.

Food-caching birds offer an opportunity to study memory in the context of navigation. These birds hide food in many distinct, concealed locations throughout their environment and later retrieve their caches. Food caching has been studied in two general types of settings well-suited for characterizing different components of the behavior. In some experiments, spatial navigation is minimized, and birds choose from a small number of available options to obtain food.^2,11–13^ These tasks have been ideal for characterizing various contributions to memory, including location, content, and even the relative time of different caches. In contrast, other experiments have been performed in the wild or in large, naturalistic settings.^14–18^ These studies have been well-suited for measuring spatial properties of the food-caching behavior, including the distribution of caches, large-scale biases, and the spatial specificity of cache memories. However, it is unknown how mnemonic and other spatial strategies are coordinated by the animal within a single behavior.

Addressing this question may be possible in reduced spatial settings, like those typical in rodent experiments.^19–21^ Such a setting must retain key spatial aspects of bird navigation, while also incorporating food-caching behavior and memory use. Ideally, it would also permit detailed tracking of the animal’s behavior and be compatible with neural recordings. We developed an experimental setup to fulfill these requirements for a food-caching species, the black-capped chickadee. We then used this setup to dissect the contributions of mnemonic and non-mnemonic navigational strategies to food caching, using probabilistic modeling of behavioral choices.^22–24^

## Results

### Design of the behavioral paradigm

Our first goal was to engineer a behavioral setup in which chickadees navigate, cache food, and retrieve caches. We designed individual cache sites as holes in the floor of an arena covered by silicone rubber flaps (Figure 1). Dimensions and materials were chosen to allow chickadees to pull the flap open with little effort – using either their toes or their beak – and to deposit a piece of a sunflower seed underneath. Once released, the flap obstructed any visual access to the contents of the site from above. Flaps ensured that no visually guided, memory-independent strategy could be used by the chickadee to determine site contents. Sixty-four cache sites were arranged in an 8×8 grid inside a 61×61 cm square arena. In addition, the arena contained a food source in each of the quadrants: a feeder with a motorized cover that could be individually opened or closed. This setup allowed navigation on a similar spatial scale to that typically studied in rodents,^20^ while also offering birds a large number of concealed cache sites.

**Figure 1.**
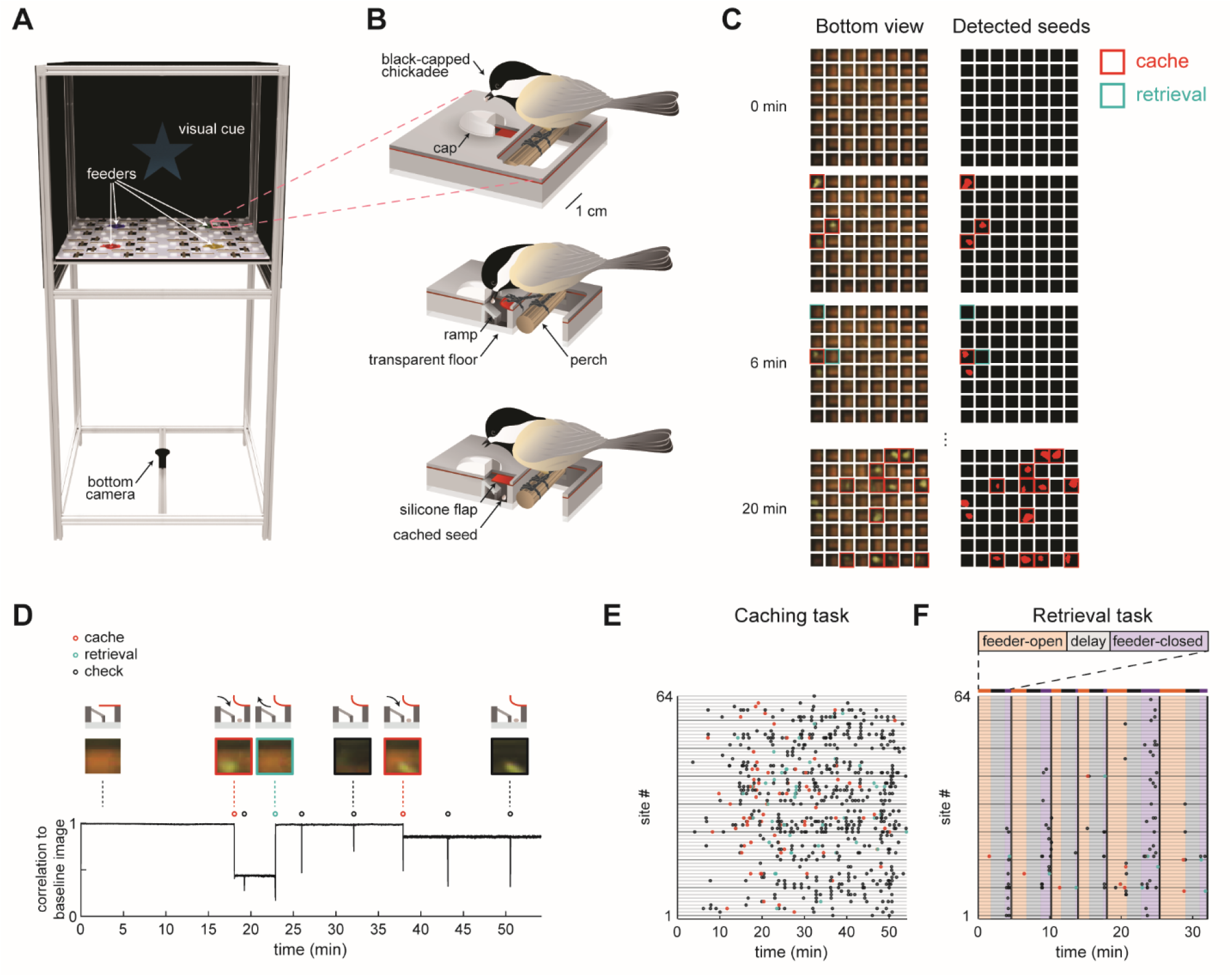
Behavioral paradigm for food caching and retrieval in chickadees. (A) Rendering of the behavioral setup. For clarity, the front wall that contains doors of the arena is removed. Pink box highlights one cache site. (B) Illustration of a chickadee at a cache site. Top: chickadee prior to caching. Middle: cross-section of the cache site, with a chickadee pulling open the silicone flap to deposit a seed. Bottom: cross-section of the cache site after the seed was cached. (C) Left: video frames from the bottom camera showing all 64 cache sites at four timepoints within a behavioral session. Right: real-time detection of cached seeds. Red area indicates shape enclosed by the detected seed contour. (D) Detection of events at one example cache site in one behavioral session. Top: cartoon of the bird’s interactions with the site at several time points. Middle: video frames from the bottom camera at the corresponding time points. Bottom: Pearson correlation of each video frame with the image of the same cache site when empty. Caches create sustained decreases in the correlation, whereas site checks create transient decreases. (E) Ethogram of all behavioral events in an example session of the Caching task. Colored circles correspond to caches, retrievals, and site checks, as in (D). (F) Same as (E), for the Retrieval task. Colored regions indicate the phase of the trial. Black vertical lines denote trial boundaries.

We next developed methods to automatically track chickadee behavior in the caching arena. Automated behavioral tracking was necessary to allow closed-loop manipulations for some of the experiments described below. To achieve this, we used a transparent material for the bottom layer of the arena and positioned a video camera underneath. We applied a real-time contour detection algorithm^25^ (see Methods) to determine whether each of the sites was empty or occupied by a seed. During offline analysis, we also detected instances of the bird opening the cover flap of a site, which we refer to as “ checks”. To do this, we calculated the Pearson correlation between the image of each site when empty and every video frame of that site in the session. Periods of time when a site was occupied corresponded to sustained periods of low correlation, while checks corresponded to transient decreases in correlation (Figure 1D). Thus, caches, retrievals, and checks could all be detected in the behavior using a single-camera video recording. We also trained a deep neural network (DeepLabCut^26,27^) to track the bird’s location for this analysis.

We designed two tasks, each suited for analyzing different aspects of behavior. In the *Caching task* (Figure 1E), chickadees were free to cache without any imposed trial structure. Individual feeders opened and closed to motivate movement through the arena and caching, but the schedule of these openings and closings was unrelated to the bird’s behavior (see Methods). Our term “ Caching task” does not mean that birds only cached in this task – in fact, birds were also free to eat seeds, check sites, and retrieve their caches. However, because this task placed no limit on the number of caches, it was particularly well-suited for investigating the chickadee’s choice of where to cache each seed.

In the *Retrieval task* (Figure 1F), a trial structure was imposed. At the beginning of each trial, one of the feeders was open (“ feeder-open phase”), allowing chickadees to eat or cache seeds. Once the bird cached 1-3 seeds, lights were turned off for a 2-min delay phase. After the delay, all feeders were closed (“ feeder-closed phase”), and the only sources of food in the arena were previously cached seeds. In the feeder-closed phase, birds were generally motivated to find and retrieve their caches. This task was therefore suitable for investigating which sites the bird chose to check during cache retrieval.

### Chickadees have unique and stable spatial biases

We first asked whether chickadees cached into some sites of the arena more than into others. In the Caching task, we calculated the cache distribution (*p*^bias^) by dividing the number of caches into each of the 64 sites across sessions by the total number of caches (Figure 2A). Birds used most of the arena for caching (between 41-64 sites for each of N=10 birds), but in many cases cache distributions appeared non-uniform. To quantify this, we measured the entropy of *p*^bias^ for each bird and compared its value to data in which caches were sampled from a uniform distribution (Figure 2B). In all birds, entropy was lower than expected by chance (p<0.001). Therefore, birds exhibited spatial biases, even though they cached widely throughout the arena.

**Figure 2.**
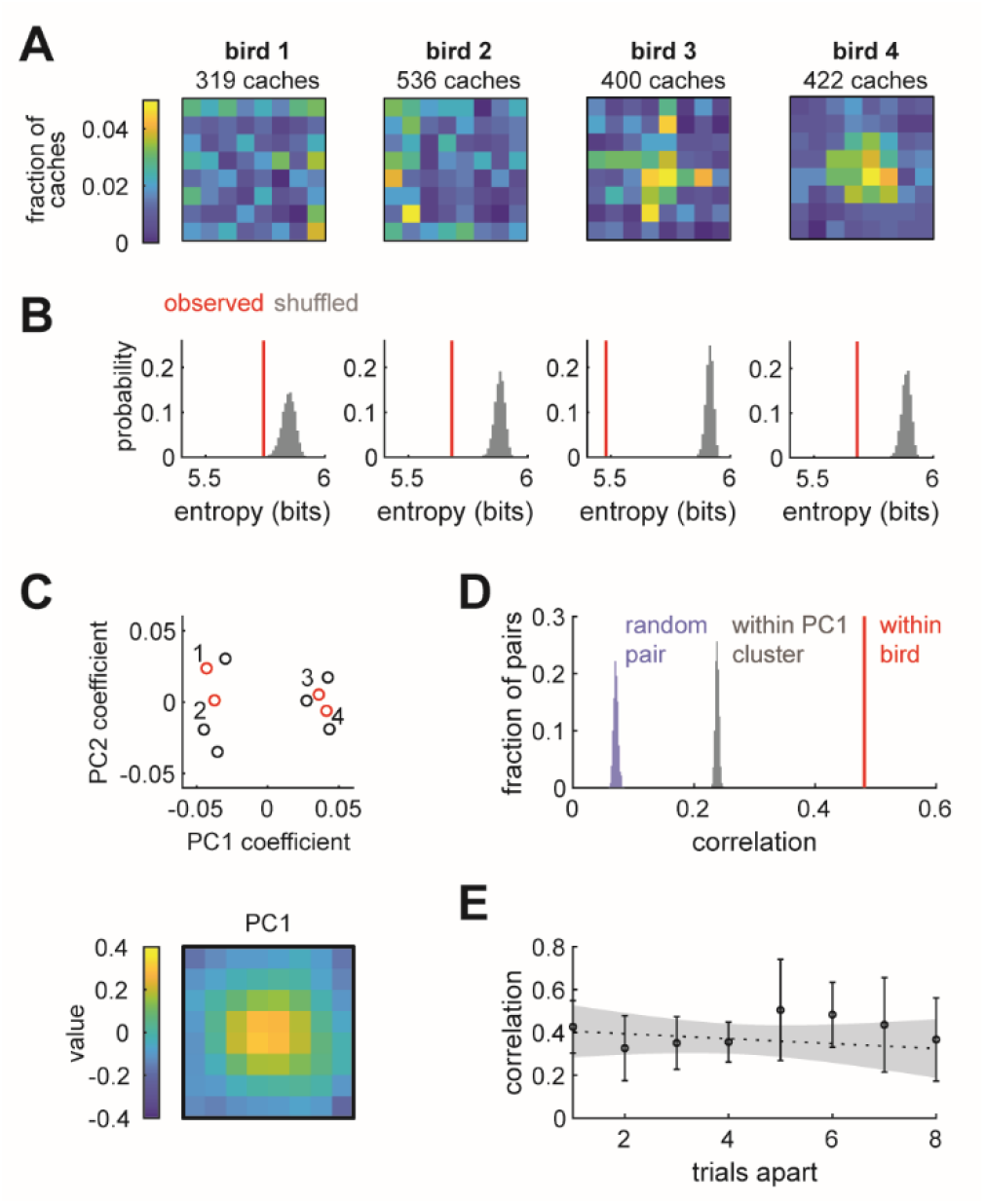
Chickadees exhibit idiosyncratic and stable biases in the locations of caches. (A) Probability distributions of cache locations across all sessions for four example birds. Distributions are denoted by p^bias^ in the text. (B) Red lines: entropy values of the spatial distributions shown in (A). Grey histograms: entropy values for simulated caches drawn from a uniform distribution. Number of simulated caches was the same as in the observed data. (C) Principal component analysis of p^bias^ values. Top: coefficients of the first two principal components for all birds. Red circles and numbers indicate birds shown in (A) and (B). Bottom: the first principal component. Birds cluster into center-preferring and edge-preferring groups. (D) Pearson correlation of p^bias^ between subsets of all sessions paired within bird, between different birds, and between birds selected from the same PC1 cluster shown in (C). (E) Pearson correlation of p^bias^ between pairs of trials at different trial lags. Vaues are medians across birds. Error bars: sem. Dashed line: linear regression. Grey shade: 95% confidence interval of linear regression.

There are several possible explanations for the observed biases. One possibility is that biases are broadly shared across individuals of the species – akin to the preference for locations near walls in rodents.^10^ Another possibility is that biases are shared across some subgroups of individuals, but not all members of the species. Finally, biases could be unique to individual birds. To distinguish between these possibilities, we first asked whether there were common types of cache distributions across birds. We calculated principal components of all *p*^bias^ values (Figure 2C). Remarkably, birds formed two clusters that were well-separated along the first principal component. The first principal component accounted for 59% of the variance across individuals and resembled a bump in the center of the arena. Thus, there were two groups of birds: those that preferred to cache in the center of the arena, and those that preferred the edges.

We next asked whether membership in one of these two groups was sufficient to explain the observed spatial biases. For each bird, we computed cache distributions separately for two randomly chosen, equal-sized subsets of the behavioral sessions. We then measured the Pearson correlation between the two distributions from the same bird (“ within-bird correlation”) and between distributions from different, randomly paired birds (“ across-bird correlation”). Within-bird correlation values were significantly higher than across-bird correlation values (Figure 2D), even when birds were paired exclusively with other birds from the same group (center-preferring or edge-preferring). Therefore, birds exhibited individual-specific biases that could not be entirely explained by common biases across a group of birds. An example of this is evident in Figure 2A: although birds 3 and 4 were both center-preferring birds, bird 3 had some additional bias for centrally-positioned rows and columns of the arena.

Finally, we asked whether spatial biases in individual birds were stable or drifted over time. We measured the Pearson correlation between cache distributions computed on pairs of sessions separated by different lags (Figure 2E). The slope of the relationship between lag and correlation was not significantly different from zero (p>0.4), indicating that there was no detectable drift in bias.

We repeated all of the above analyses for the distribution of site checks in the Retrieval task (N=7 birds). In this case, *p*^bias^ was computed by dividing the number of times a particular site was checked by the total number of all site checks. All of the results were similar to the ones obtained for caches in the Caching task (Figure S1). Collectively, our results suggest that chickadees exhibit idiosyncratic, but stable biases, both in the sites that they check and those that they choose for caching. In both cases, the *p*^bias^ distribution therefore serves as a valid baseline for subsequent models, described below.

### Probabilistic models of chickadee behaviors

How do chickadees choose where to cache? One model consistent with our analysis so far is that birds choose each cache location by drawing it randomly from the distribution *p*^bias^. However, this is not the only possibility because *p*^bias^ is computed by pooling across all time points. On a moment-by-moment basis, the likelihood of caching into a particular site may differ from the value given by *p*^bias^. For example, the choice of cache site could depend on the proximity of a site to the bird: chickadees may tend to cache close to the location they last interacted with or, conversely, they may avoid the vicinity of that location. The choice of a cache site may also depend on which sites are already occupied by seeds, since birds may tend to either cluster or to spread out their caches. The bird’s knowledge of which sites are occupied might be updated either by the act of caching, or by the act of checking the site to view its contents.

To model these possibilities, we considered three types of “ special” places in the arena at each moment during the Caching task: 1) the location of the last site that the bird interacted with, which includes checking and/or retrieving from one of the sites, or visiting one of the feeders, 2) sites that are currently occupied by seeds, and 3) sites that have been previously checked by the bird and are currently empty (“ checked-empty” sites). We then constructed three scaling factors whose values changed with proximity to each of these special places (Figure 3, defined in Methods). The likelihood of caching into a particular site was computed by multiplying all three scaling factors by the corresponding value of *p*^bias^ and normalizing the product to a sum of 1 across the arena. Each of the scaling factors was defined by two parameters. The first parameter (*γ*_prv_, *γ*_occ_, or *γ*_emp_ for the previous interaction, occupied sites, and checked-empty sites, respectively) indicated the strength and direction of the corresponding effect on probability, measured on a logarithmic base-10 scale. For example, *γ*_occ_ =−1meant that the act of caching into a site decreased the likelihood of caching there again by a factor of 10. The second parameter (σ_prv_, σ_occ_, or σ_emp_, respectively) indicated the spatial extent of each effect. In other words, a large value of σ_occ_ meant that the act of caching into a site affected the likelihood of subsequently caching into neighboring sites as well. This model was first applied to the data obtained in the Caching task. For each bird, we optimized the values of the model parameters in order to maximize the likelihood of the experimentally observed caches.^22–24^

**Figure 3.**
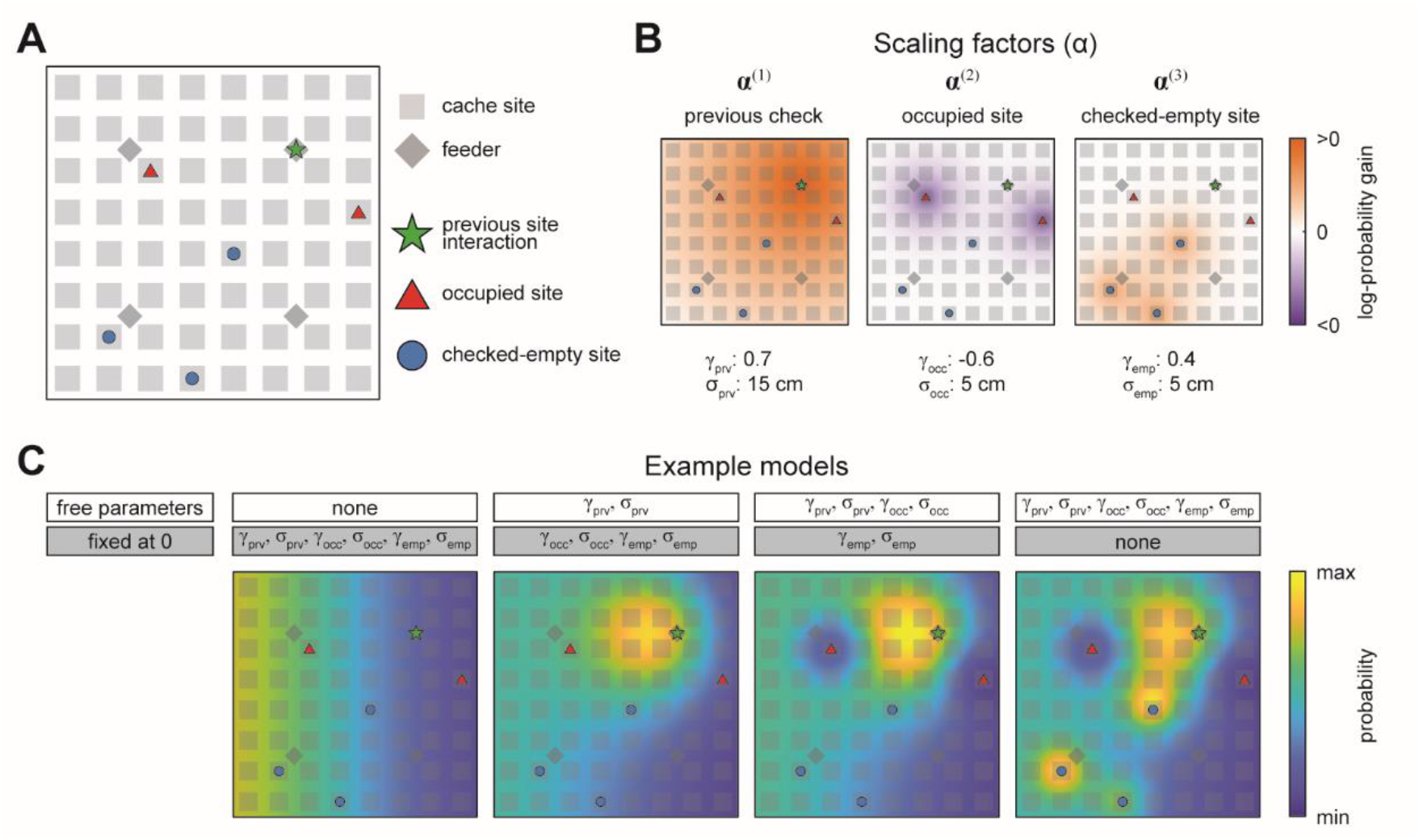
Schematic of the probabilistic behavioral models. Models here compute the probability of caching into a particular site in the Caching task. Equivalent models are used for the probability of checking a particular site in the Retrieval task. (A) Schematic of the arena at an example time point in the Caching task depicting “ special locations” in the arena: previous site interaction, occupied sites, and checked-empty sites. (B) Examples of scaling factors that change with proximity to each of the special locations shown in (A). Probability distribution of caching into different sites is computed by multiplying all scaling factors by p^bias^. (C) Example models. For each model, some parameters are fixed at 0, while others are free and fit using maximum-likelihood estimation. For free parameters, examples shown here use the same values as in (B). All four models are plotted on the same color scale.

We also asked how chickadees chose which sites to check in the Retrieval task. Much like the choice of cache location, the choice of check location may be affected by proximity to the previously checked site, to occupied sites, and to checked-empty sites. We therefore used the same model as above, but fit to locations of checks instead of caches. At first, we fit the model only to the checks that the bird made up until and including finding a cache in the Retrieval task. We did this because after finding the first cache, birds often recached food, which confounded the analysis by mixing caching and retrieval behaviors in the same time period. We analyze recaching separately below.

Our approach was to start with a baseline model (Model #0), in which the likelihood of choosing a particular site for caching or checking was given by *p*^bias^. In this model, there were no free parameters, and the values of *γ*_prv_, *γ*_occ_, *γ*_emp_, σ_prv_, σ_occ_, and σ_emp_ were all set to 0. We then gradually introduced free parameters to the model, one or two at a time. With each parameter introduction, we tested whether model performance was improved by measuring the Akaike Information Criterion (AIC), which quantifies model likelihood accounting for the number of free parameters.^28,29^

### Chickadee behaviors exhibit a strong proximity effect

We asked how the location of the previous site interaction affected the chickadee’s behavior. For this analysis we constructed Model #1, in which *γ*_prv_ and σ_prv_ were introduced as free parameters (Figure 4). We found that the AIC for Model #1 was lower than for Model #0 (p<0.001), indicating that these two parameters significantly improved model performance. The best-fit value of *γ*_prv_ was positive in all birds (0.94 ± 0.21, p<0.001), and the best-fit value of σ_prv_ was 15.5 ± 0.9 cm. Thus, there was roughly a 10-fold (i.e., 10^0.94^) increase in probability of caching next to the previous site interaction, and the spatial extent of this effect was about 1/4 of the arena. To corroborate this result independently of the model, we compared the observed and expected distances of caches to previous site interactions and indeed found that birds cached closer to previous sites than expected by chance (Figure S2).

**Figure 4.**
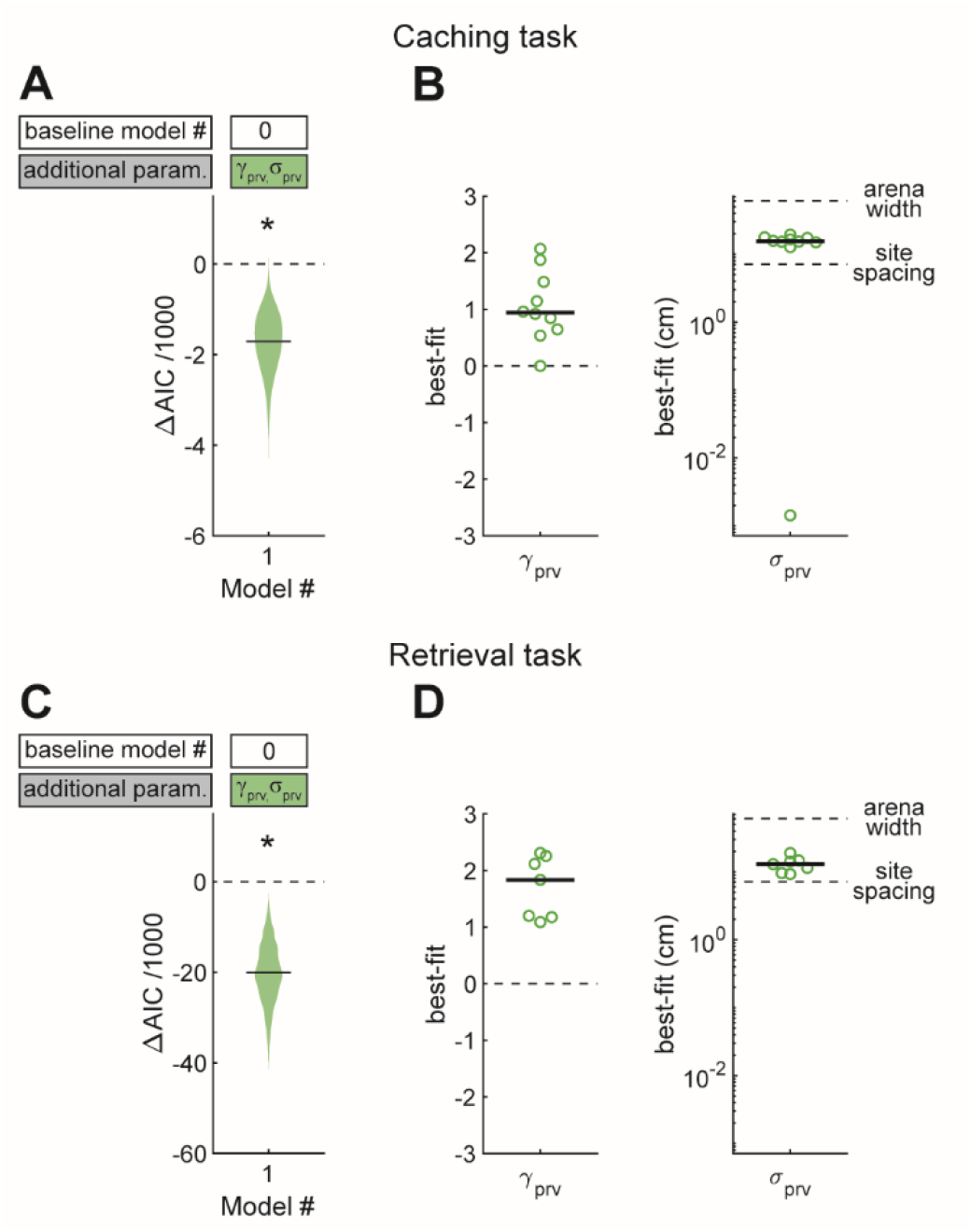
Effect of proximity to the previous site in Caching and Retrieval tasks. (A) Comparison of Model #1 to Model #0 applied to the Caching task. Here and in subsequent figures, ΔAIC indicates the difference in AIC between the model being considered (in this case Model #1, labeled on the x-axis) and the baseline model (in this case Model #0, labeled in the white box). Compared to Model #0, Model #1 uses two additional free parameters (*γ*_prv_ and σ_prv_), specified in the colored box. Horizontal black line: ΔAIC value for caches pooled from all birds. Shaded area: distribution of 1000 ΔAIC values on data bootstrapped across birds. Values less than 0 indicate model improvement. Asterisk indicates statistically significant improvement. (B) Best-fit values of model parameters *γ*_prv_ and σ_prv_ for Model #1 applied to the Caching task. Symbols indicate values for individual birds. Black line: median values across birds. (C, D) Same as (A, B), but for models applied to the Retrieval task.

We applied the same model to site checks in the Retrieval task. Again, AIC for Model #1 was lower than for Model #0 (p<0.001). The value of *γ*_prv_ was positive in all birds (1.83 ± 0.42, p<0.001), indicating a tendency to check sites close to the previous site interaction, and σ_prv_ was 12.9 ± 1.7 cm. This result is consistent with our qualitative observations of chickadees in the Retrieval task: as birds moved through the arena – sometimes in the direction of a hidden cache –they often checked sites on their path.

### Chickadee behaviors are affected by site content

We next asked whether chickadee behaviors were additionally influenced by the contents of individual sites (Figure 5A-B). In the Caching task, we fit Model #2, which included *γ*_occ_ as a third free parameter to model the effect of occupied sites. The AIC value for Model #2 was lower than for Model #1 (p<0.001), indicating that occupied sites indeed affected the behavior. We then fit Model #3, in which *γ*_emp_ was additionally introduced as a fourth free parameter to model the effect of checked-empty sites. Model #3 had an even lower value of AIC (p<0.001), indicating that checked-empty sites also affected the behavior. Across birds, the value of *γ*_occ_ was negative (−0.32 ± 0.17, p<0.002), whereas the value of *γ*_emp_ was positive (0.13 ± 0.05, p<0.001). These values indicate roughly a 3-fold decrease in probability of caching into occupied sites and a 1.3-fold increase in probability of caching into checked-empty sites. Both results are consistent with chickadees spreading their caches throughout the arena, rather than pooling them into the same sites. Indeed, chickadees tended to cache into occupied sites less often than expected by chance (Figure S3A).

**Figure 5.**
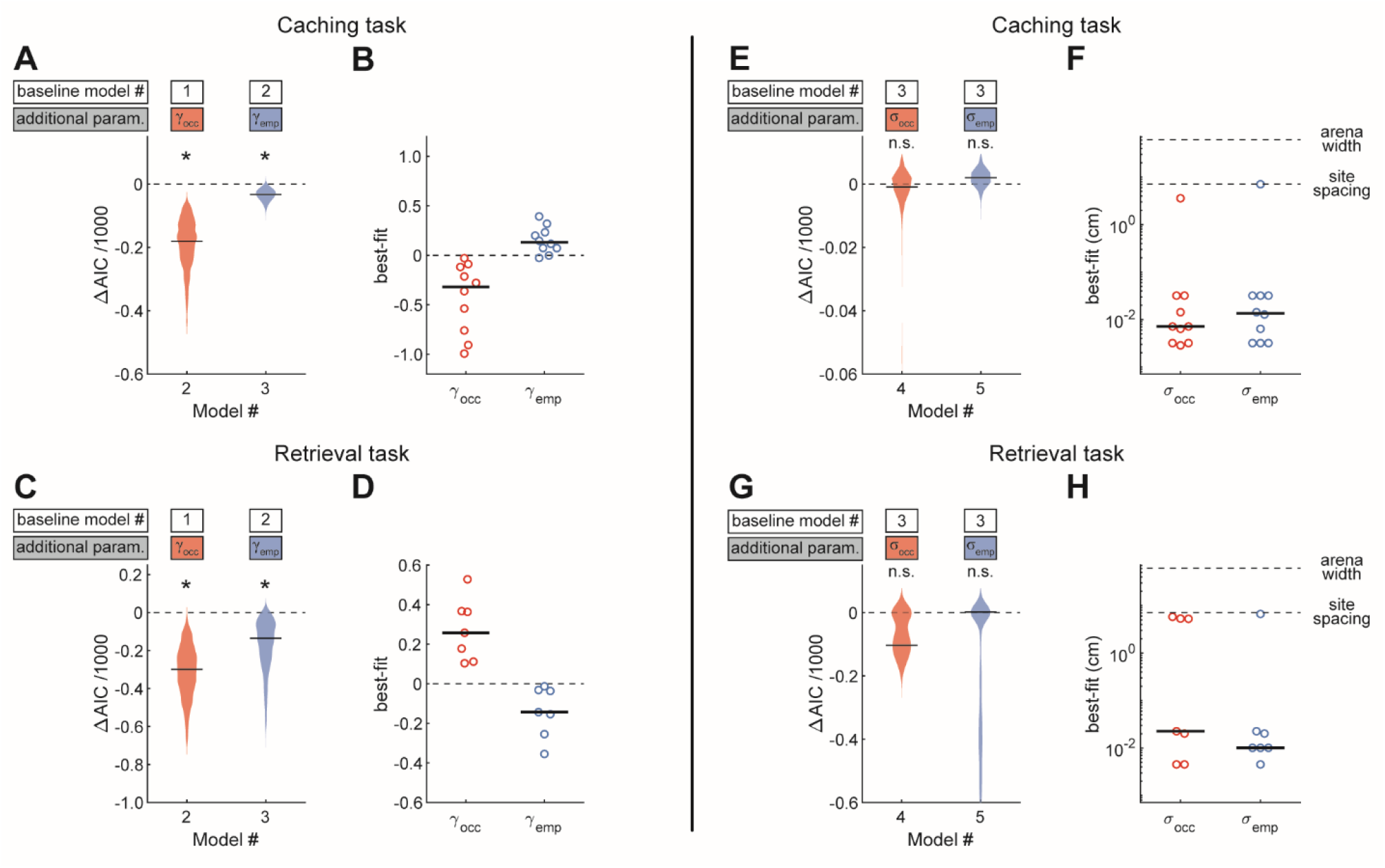
Site-specific, opposing effects of site content on behavior in Caching and Retrieving tasks. (A) Performance of Models #2 and #3 applied to the Caching task, plotted as in Figure 4A. These models introduce free parameters to quantify the effect of occupied and checked-empty sites on behavior. (B) Best-fit values of *γ*_occ_ and *γ*_emp_ applied to the Caching task, plotted as in Figure 4B. (C, D) Same as (A, B) but for models applied to the Retrieval task. (E) Performance of Models #4 and #5 applied to the Caching task, plotted as in Figure 4A. These models introduce free parameters to quantify the spatial extent of the effect of occupied and checked-empty sites on behavior. (F) Best-fit values of σ_occ_ and σ_emp_ applied to individual birds in the Caching task, plotted as in Figure 4B. (G, H) Same as (E, F), but for models applied to the Retrieval task.

We again applied the same models to site checks in the Retrieval task (Figure 5C-D). Like in the Caching task, introducing *γ*_occ_ and *γ*_emp_ significantly improved the fit of the model (change in AIC<0, p<0.001 and p<0.02, respectively). However, the signs of the two parameters were reversed compared to the Caching task: *γ*_occ_ was positive in all birds (0.26 ± 0.09, p<0.005), whereas *γ*_emp_ was negative in all birds (−0.14 ± 0.07, p<0.03). These results show that birds were attracted to occupied sites during the feeder-closed phase of the task. This is consistent with chickadees searching for their caches. Indeed, chickadees found their caches after fewer checks than expected by chance: 8.5 ± 5.6 checks in the observed data, compared to 20.8 ± 4.5 checks in data where trajectories were shuffled between trials (Figure S3B).

Our analyses so far show that chickadees can be attracted to or avoid specific sites depending on the content of these sites. We asked whether these behavioral effects were site-specific, or whether neighboring locations were affected as well. We introduced either σ_occ_ or σ_emp_ as an additional free parameter (Model #4 and Model #5 respectively, Figure 5E-H). In both tasks, the AIC values for Model #4 and Model #5 were not significantly lower than for Model #3. Therefore, neither σ_occ_ nor σ_emp_ improved the fit of the model. In fact, the best-fit values of σ_occ_ and σ_emp_ in both models were between 0.01-0.02 cm, much smaller than the distance between neighboring sites (7.1 cm). Thus, the effects of site contents on chickadee behavior were site-specific, and were small even at neighboring sites of the arena. We corroborated this result independently of the model by measuring the fraction of caches and checks at various distances away from occupied sites (Figure S3C-D).

### Chickadees have memories of site contents

We have shown that chickadees avoid caching into sites that are occupied, but increase their probability of caching into checked-empty sites. An intriguing explanation of these results is that chickadees use memory of site contents to influence their caching behavior. However, after obtaining a seed, a chickadee may simply check multiple sites in the arena and preferentially cache into an empty one that it encounters. This would be a visually-guided strategy that does not necessarily require memory. We considered this possibility unlikely because chickadees usually cached into the first site that they opened after visiting the food source (79% of caches). However, we wanted to ensure that the remaining 21% of caches were not responsible for the observed effects. We implemented the same models as before (Models #2 and #3), but instead of fitting them to the observed caches, we fit them to the locations of the first sites opened by the bird after obtaining a seed (“ first check”). We found that *γ*_occ_ in Model #2 was still significantly negative, and *γ*_emp_ in Model #3 was still significantly positive (Figure 6A-B, p<0.005 and p<0.03, respectively). Thus, after a chickadee obtained a seed for caching, the very first site it opened had an increased probability of being empty. This result implies that, during caching, chickadees used memories of which sites were occupied and which were empty.

**Figure 6.**
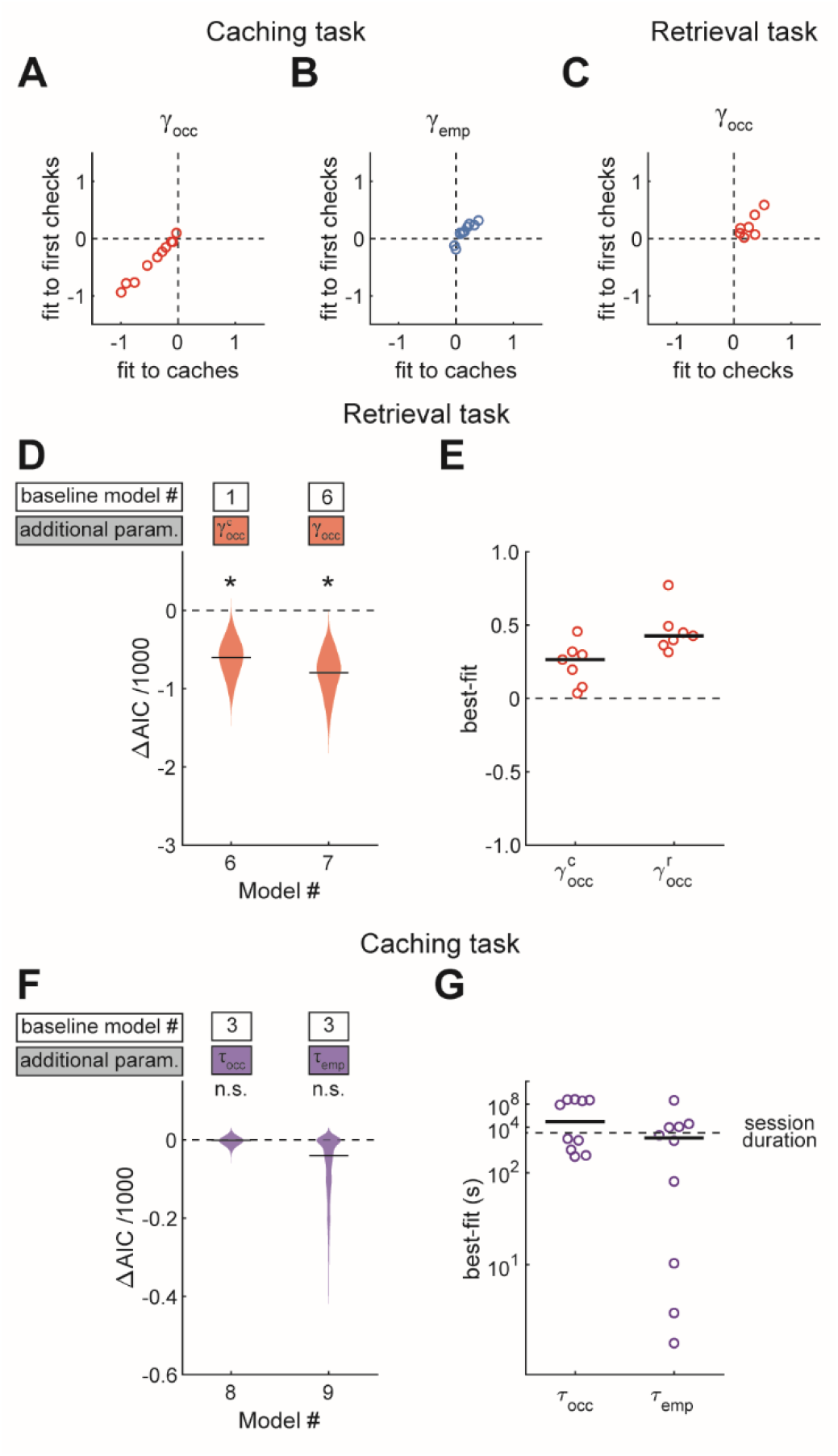
Results are best explained by long-lasting memories. (A, B) Values of parameters across birds, compared between models applied to caches or first checks in the Caching task. Values of *γ*_occ_ are from Model #2, and values of *γ*_emp_ are from Model #3. (C) Values of *γ*_occ_ across birds, compared between Model #2 applied to checks or the same model applied to first checks in the Retrieval task. (D) Performance of Models #6 and #7 applied to the Retrieval task, plotted as in Figure 4A. These models introduce free parameters to separately quantify the effects of caches and recaches on the site checking behavior. (E) Best-fit values of 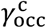 and 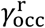 applied to individual birds in the Retrieval task, plotted as in Figure 4B. (F) Performance of Models #8 and #9 applied to the Caching task, plotted as in Figure 4A. These models introduce free parameters τ_occ_ and τ_emp_ to quantify the decay of *γ*_occ_ and *γ*_emp_ over time since the site was last checked by the bird. (G) Best-fit values of τ_occ_ and τ_emp_ applied to individual birds in the Caching task, plotted as in Figure 4B.

We also fit Model #2 to first checks in the Retrieval task. In this case, *γ*_occ_ was still positive (p<0.02). In other words, chickadees had an increased probability of checking an occupied site even on the first attempt. This implies that chickadees did not search the arena until finding a cache, but used memories of occupied sites. Note that *γ*_emp_ was not computed for this analysis because, by definition, there could not be any checked-empty sites before the first check.

Finally, we wanted to make sure that chickadees did not use olfactory cues to detect seeds in cache sites. Chickadees are not known to use olfaction for finding food,^14,15,18^ and previous studies have excluded this possibility by removing caches during the delay period of the task. We repeated this experiment in our arena. Consistent with published results, even when caches were removed from the arena, chickadees tended to spend more time at the locations of the missing caches after the delay period (Figure S4). Collectively, our results show that chickadees have memories of site contents and use them in flexible ways, both during caching and during retrieval.

### Memories of site contents are long-lasting

We next asked how long memories lasted in our arena. In the feeder-closed phase of the Retrieval task, chickadees often “ recached” seeds after retrieving them (44% of all retrievals). Therefore, there were two types of caches in this task: recent recaches and older caches made prior to the delay period. We first analyzed how these different types of caches affected behavior. For this analysis, we introduced models with separate parameters 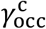 and 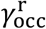 instead of 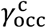 to fit the contribution of older caches and recaches, respectively. These models were fit to the entire feeder-closed phase of the Retrieval task, which contained both the retrieval and the recaching behaviors. Model #6 introduced 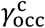 as a free parameter, while Model #7 introduced both 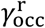 and 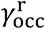. We found that Model #6 fit the data significantly better than Model #1, and Model #7 fit the data even better than Model #6 (Figure 6D, change in AIC<0, p<0.001 in both cases). The best-fit values were significantly positive for both parameters, but higher for 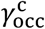 than for 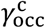 (0.42 ± 0.05 and 0.26 ± 0.07, respectively, p<0.01). These results indicate that birds were attracted to all of their caches during retrieval, but had a stronger preference for the recent recaches.

One possible explanation of this result is that birds had a memory decay that reduced the effect of older caches on behavior. However, birds often recache a single seed multiple times, and may simply have a preference for revisiting recache locations. We therefore performed a different analysis that modeled a continuous memory decay. The Caching task was ideally suited for this analysis because it did not have a trial structure; caches could be retrieved at any later time in the session. We defined a “ memory decay factor” for each site, which was reset to 1 whenever that site was checked and decayed exponentially to 0 afterwards. In Model #8, this factor was multiplied by *γ*_occ_ and decayed with a timescale *τ*_occ_. In Model #9, this factor was multiplied by *γ*_emp_ and decayed with a timescale *τ*_emp_. Both models were compared to Model #3, in which neither *γ*_occ_ nor *γ*_emp_ decayed. We found that neither Model #8 nor Model #9 improved the fit to the data (change in AIC not significantly negative, Figure 6F). The best-fit values of *τ*_occ_ and *τ*_emp_ were very large (both >44 min, Figure 6G) compared to the median time between consecutive checks of the same site (2.4 min). Therefore, there did not appear to be a significant memory decay at the timescale of our experiments. This additionally implies that the preference for retrieving recaches may be a behavioral preference unrelated to a memory decay.

### Memories of site contents are high-capacity and accessed in an arbitrary order

Food-caching birds are famous for storing and likely remembering large quantities of food items in the wild.^30^ So far, it is unclear if chickadees exhibit a high memory capacity in our arena. For example, the statistical effects we observed can be explained by chickadees remembering a small number of initial caches and having no memory of the remaining caches. To test this possibility, we allowed *γ*_occ_ and *γ*_emp_ to decay exponentially with the total number of occupied sites and checked-empty sites in the arena. The decay constants of these exponentials were two additional parameters, *v*_occ_ and *v*_emp_, respectively. As before, we fit two models (Model #10 and Model #11), each introducing one of these two parameters. Neither model fit the data better than Model #3, in which *γ*_occ_ and *γ*_emp_ did not decay (change in AIC not significantly negative, Figure 7A). The best-fit values of *v*_occ_ and *v*_emp_ were over 18 sites in both cases (Figure 7B) – large compared even to the maximum number caches present in the arena at any single moment in the session (15.0 ± 1.0, N = 101 sessions). Therefore, memory did not appear to decay at capacities tested by our experiments.

**Figure 7.**
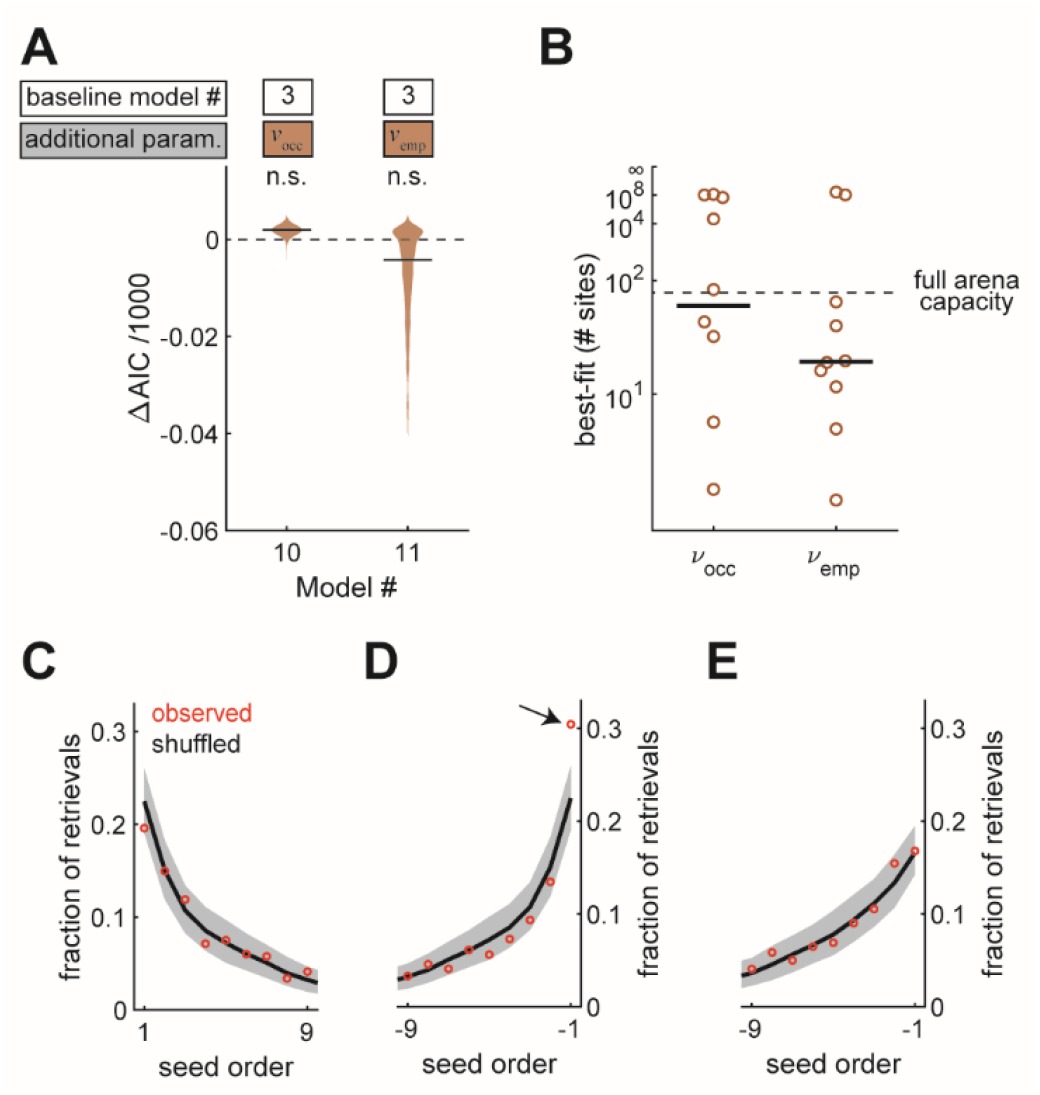
Memories are high capacity, and accessed in arbitrary order. (A) Performance of Models #10 and #11 applied to the Caching task, plotted as in Figure 4A. These models introduce free parameters *ν*_occ_ and *ν*_emp_ to quantify the decay of *γ*_occ_ and *γ*_emp_ as either the number of occupied sites or the number of checked-empty sites in the arena increases, respectively. (B) Best-fit values of *ν*_occ_ and *ν*_emp_ applied to individual birds in the Caching task, plotted as in Figure 4B. (C) Fraction of retrievals as a function of order in which seeds were caches. Order is aligned to the first cache. Red marker: mean observed fraction across birds. Black line: average value expected from retrieving seeds in a random order. Grey: 95% confidence interval of retrievals made in a random order. (D) Same as (C), but aligned to the most recent cache. Arrow indicates the only point that was significantly outside the 95% confidence interval of shuffled data. (E) Same as (D), but with transient caches eliminated. Transient caches are those that are retrieved without leaving the perch.

All of our models so far have used spatial location to explain the choice of sites for caching and checking. We also asked whether the order in which seeds were cached was predictive of the order in which birds chose to retrieve them. For this analysis, we used the Caching task, in which birds often cached and retrieved large numbers of seeds. Across all retrievals, we asked how often the *n*th cached seed was chosen by the bird (Figure 7C). There was no difference between observed data and shuffled data, in which seeds were chosen randomly from the available caches. Thus, there was no detectable “ primacy” effect in the data. We then asked how often the *n*th most recent seed was chosen. In this case, the most recently cached seed was retrieved more often than expected by chance (p<0.001, Figure 7D). However, this “ recency” effect was largely caused by caches that the bird retrieved without leaving the site of the cache (Figure 7E). We often observed chickadees making several of these “ transient caches” and retrievals with a single seed, before finally settling on a cache location. Once transient caches were excluded from the data, there was no detectable primacy or recency effect in the order of retrieval.

## Discussion

In this study, we designed an apparatus to explore how chickadees make navigational decisions during food caching and retrieval. We built quantitative models to understand the relative contributions of mnemonic and non-mnemonic strategies to these behaviors. Our setup allowed birds to move within an arena similar to those used for rodent experiments.^21,31,32^ We found that several natural features of chickadee spatial navigation were preserved in this environment. Our apparatus also allowed birds to cache and retrieve food from a relatively large number of concealed sites within the arena. Tracking behavior at these sites revealed that chickadees rely on multiple strategies for caching and retrieval and use memory in surprisingly flexible ways during these behaviors.

We found two spatial strategies that did not require birds to remember the contents of individual sites. First, chickadees had spatial biases that were stable over time and unique to individual birds. Previous work has shown spatial biases in the wild. For example, individual birds can cache in different zones within a shared territory or on different parts of a tree.^9,14,33^ This strategy has a clear ethological advantage: biases reduce the memory load of cache locations, and choosing biases idiosyncratically minimizes unwanted overlap between individuals. The second contribution to behavior was the effect of proximity: chickadees tended to cache and check sites close to their previous site interaction. These effects have also been observed in the wild and have ethological advantages. Caching close to the food source minimizes travel distance and exposure to dangers.^34–36^ Checking locations in close proximity is an efficient opportunistic method of exploring the environment.^7,37^ Although the patterns we find are on a dramatically smaller scale, it is intriguing to speculate that biases and proximity effects are driven by shared mechanisms across conditions.

We do not know whether these effects are truly non-mnemonic on timescales longer than our experiment. For example, a chickadee’s experiences over long periods of time may influence their biases or their typical trajectories within an environment.^38^ However, for the purposes of our experiment, explaining away biases and proximity effects was critical for isolating the remaining memory-guided components.

Our analysis has revealed several features of chickadee memory. Memories were spatially specific, long-lasting, high-capacity, and could be retrieved in an order different from their storage order. These features have been studied before – usually in experiments specifically designed to look at each feature of memory in isolation. For example, radioactive tagging of food in the wild has shown that birds search for caches with centimeter precision.^14^ In both field and laboratory experiments, birds continue to remember locations at least a month later.^39,40^ Chickadees can cache thousands of seeds per day and are suspected to remember a large fraction of them.^30,41^ Finally, previous studies have also not detected a relationship between the order of caching and the order of retrieval, suggesting that chickadees have independent memories of caches rather than a single memory of the action sequence.^15,42,43^ Our models capture these features on smaller spatial and temporal scales, and notably combine them within a single laboratory paradigm.

Classic work has shown that food-caching birds use memory to retrieve food.^11,15,16^ They can furthermore remember the contents of individual sites – for example, to choose caches that contain a certain type of food.^11,15^ We also observed memories of site contents in our arena. Chickadees were attracted to occupied sites during retrieval, but avoided empty sites that they had previously checked. These results show that memories of site contents in chickadees not only affect a single behavior, but even influence the choice between different behaviors – in our case, site attraction or avoidance.

Site content influenced not only retrieval, but also caching itself. When caching, birds avoided occupied sites, but preferred sites that they had checked and found to be empty. These behaviors resulted in a “ spreading” of caches throughout an environment. Cache spreading is observed on a larger scale in the wild and is thought to be a defensive strategy against pilferers.^14,44^ However, it was not previously known that memory plays a role in this behavior. The implication of our results is that a single memory of a cached seed can be used by the bird for entirely different behaviors depending on context – in our case, caching or retrieval. The ability of the same memory to flexibly drive different behaviors is a hallmark of episodic memories in humans.^19,45–47^ It has been suggested that cache memory in birds is similarly flexible, but this idea has remained controversial.^48,49^ Our finding that the memories can be used for different goals, both in a content- and context-dependent way, lends support to this idea.

In summary, our behavioral apparatus brings a complex, natural behavior into the constraints of a laboratory setup. In this reduced preparation, behavior is automated and highly trackable, allowing in-depth analysis and closed-loop experimental manipulations. Notably, by being relatively small, flat, and unobstructed, this arena is compatible with a wide array of techniques, such as tethered recordings.^50^ Our behavioral paradigm therefore offers a powerful tool for investigating the interaction of episodic-like memories with spatial behaviors.

## Acknowledgements

This work was supported by the NSF Graduate Research Fellowship Program (MCA), NIH training grant T32 EY013933 (MCA), New York Stem Cell Foundation – Robertson Neuroscience Investigator Award, and the Beckman Young Investigator Award. We thank D. Scheck, S. Hale, T. Tabachnik, R. Hormigo, and K. Gutnichenko for technical assistance; the Black Rock Forest Consortium, J. Scribner and the Hickory Hill Farm, and T. Green for help with field work; L. Abbott and members of the Aronov lab for comments on the manuscript. Illustrations in Figure 1B are by J. Kuhl.

## Author contributions

M.C.A. and D.A. conceived of the experiments. M.C.A. performed the experiments. M.C.A. and D.A. analyzed data and wrote the manuscript.

## STAR Methods

### Resource availability

#### Lead contact

Further information and requests for resources and reagents should be directed to and will be fulfilled by the Lead Contact, Dmitriy Aronov.

#### Materials availability

This study did not generate new unique reagents.

#### Data and code availability

Scripts used for behavioral analysis will be available on GitHub following publication. Requests for further details of the software and raw data should be directed to and will be fulfilled by the Lead Contact, Dmitriy Aronov

### Experimental model and subject details

All animal procedures were approved by the Columbia University Institutional Animal Care and Use Committee and carried out in accordance with the US National Institutes of Health guidelines. The subjects were 21 adult black-capped chickadees (*Poecile atricapillus*) collected from multiple sites in New York State using Federal and State scientific collection licenses. Subjects were at least 4 months old at the time of the experiment, but age was not determined more precisely. Chickadees are not visibly sexually dimorphic, and all experiments were performed blindly to sex. Sex was only determined on birds that were subsequently used for other procedures in the lab. For the Caching task, 17 chickadees were used (7 male, 3 female, 7 unknown). Of these birds, 7 made fewer than 64 caches and were excluded from all analyses except those in Figure S3A,C. For the Retrieval task, 7 chickadees were used, 6 of which had been previously run on the Caching task (3 male, 1 female, 3 unknown). The control experiment described in Figure S1 was performed on 3 birds, none of which were used in other tasks (sex unknown). Prior to experiments, all birds were housed in groups of 1-4 on a “ winter” light cycle (9h:15h light:dark).

### Behavioral apparatus

#### Design of caching apparatus

All custom parts of the behavioral arena were designed in Autodesk Inventor. Part files are available upon request. Two identical enclosed square arenas were constructed and used for running all experiments. Each arena was a 61 cm x 61 cm square and contained 64 perches, 64 “ cache sites”, and 4 “ feeder sites”.

The arena was constructed from five laser-cut layers. The top surface layer was a white polystyrene sheet (1.6 mm thick, McMaster-Carr, 8734K32). The second layer was a sheet of silicone rubber (0.8 mm thick, 60A durometer, McMaster-Carr 1460N41). The third and fourth layers were black cast acrylic sheets (3.2 mm and 4.8 mm thick, McMaster-Carr, 8505K742; 8505K748, respectively). Finally, the fifth (bottom-most) layer of the arena was a transparent cast acrylic sheet (3.2 mm thick, McMaster-Carr 8560K257).

All five layers contained cutouts for the perches. These included space directly in front of and behind the perch, allowing birds to wrap their toes around the perch. The fourth layer included small ledges on which the ends of the perch rested. Perches were 9.5 mm diameter, 50.8 mm length wooden dowels (McMaster-Carr, 97195A434).

Cache sites were created by making square cutouts in the third and fourth layer. These cutouts formed a cavity 6.4 mm wide x 4.8 mm long x 7.9 mm deep. A 3D-printed ramp was inserted into this cavity and helped direct seeds into the bottom of the site (Clear Resin, Formlabs, RS-F2-GPCL-04). The bottom (fifth) layer contained no cutouts at the cache sites and therefore provided a floor for seeds to rest on. The second (silicone rubber) layer was cut on three sides above each cache site, creating a soft flap that could be pulled open. The top layer had a cutout above each site, providing access to the silicone flap. Finally, a white 3D-printed cap was attached to the top level next to each site (White Resin, Formlabs RS-F2-GPWH-04). This cap created a barrier that blocked visual access of site contents from anywhere in the arena.

The top four layers were screwed together. The bottom layer was attached using small magnets (K&J Magnetics, D22), and could be temporarily removed in order to clean out all cache sites between sessions.

Each feeder consisted of a 3D-printed basket (White Resin, Formlabs RS-F2-GPWH-04) attached to a stepper motor (Mouser, 108990003) that could open or close the feeder. The motor was controlled by an Arduino (Mouser, 713-102990189) via a stepper motor driver (Mouser, 474-ROB-12779). A ledge surrounding each feeder was 3D printed from colored PLA using a LulzBot Taz 6 3D printer. Each feeder’s ledge was a different color (red, blue, green, and yellow). Lights in the arena were turned on and off using a solid-state relay (Mouser, 558-D1D12), also controlled via the Arduino.

Each arena was placed inside a black plastic box with doors on one side. Walls of the box were positioned 2.5 cm from the edge of an arena, creating a “ moat” that almost completely surrounded the arena (except the corners). The box was 51 high. Bright shapes ∼15 cm across (blue star, orange triangle, purple circle, and green tree) were positioned on the center of each wall. The arena was illuminated both from above and from below the environment, using LED lights (Super Bright LEDs, VTL -x1515; NFLS-WW300X2-LC2). Natural sound of rushing water was played in the background to mask inadvertent room noises.

Birds were monitored using three cameras. A camera in the center of the ceiling was used to track bird’s position (Amazon, 180-degree Fisheye lens 1080p Wide Angle Pc Web USB camera, 30 fps). A camera positioned 90 cm below the arena (bottom camera) monitored the contents of cache sites and the bird’s site checks (Edmund Optics, acA2500-60uc; #63-245, 1” PYTHON 5000 CMOS sensor, 57 fps). A third camera mounted in the center of one wall 24 cm above the floor of the arena (side camera) and was used for manual verification of bird’s behavior (Edmund Optics, EO-2323; #62-274, 2/3” Progressive Scan CMOS sensor, 50 fps).

#### Actuation of behavioral arena

For the Caching task, where no closed-loop manipulation was needed, all actuation (control of lights and motors) was performed by standalone Arduino code. For the Retrieval task, all task-relevant information (contents of cache sites and the bird’s location) were monitored in real time by software written in Bonsai.^25^ This software communicated with the Arduino to actuate events in the arena. Bonsai received real-time inputs from the bottom camera and the top camera. All code is available upon request.

To detect occupied sites, an ROI was drawn around each cache site in the green channel of video recorded by the bottom camera. The green channel was used because it provided high contrast with the red silicone material. At the beginning of each trial, the background of each ROI was subtracted using *BackgroundSubtraction* (subtractionMethod Absolute, ThresholdType ToZero, ThresholdValue 6). In each frame, novel objects were then detected using *FindContours* (Method ChainApproxNone, minArea 4) and counted using *BinaryRegionAnalysis*. The count of occupied sites was then passed through a lowpass filter (y_*n*_ = 0.01*x*_*n*_ + 0.99*y*_*n*−1_, where *x*_*n*_ is the count and *y*_*n*_ is the filtered count). The value of y_*n*_ rounded to the nearest integer was used by Bonsai as the current number of seeds in the arena. This filter introduced a delay of ∼1 s between a cache occurring and being detected, ensuring that checks were not counted as caches. An experimenter was present during all behavioral sessions to verify the performance of the automated tracking.

To detect a visit to a feeder, an ROI was drawn around each feeder in video recorded by the top camera. The sum of pixel values in this ROI was computed, and a threshold was chosen separately for each feeder to reliably detect the chickadee’s head entering the ROI. Because chickadees are black-capped, all feeders were lighter in color, and ROI entrances corresponded to negative threshold crossing.

### Behavioral protocol

#### Habituation

Birds selected for experiments were singly housed and weighed daily. Primary feathers were trimmed to prevent flight. Initially, birds were given an ad-libitum supply of Mazuri Small Bird Diet. For some birds, the diet was supplemented with dried mealworms. Each bird’s cage contained a small platform with 4-6 “ replica” cache sites identical to those later used in the behavioral arena.

Birds were first given a 4-day period of acclimation and habituation to handling. After 4 days, they were gradually habituated to food restriction. First, birds were not given food for the first two hours of the lights-<u>on</u> period. This period was gradually increased to 4 hours (occasionally 3-5). Birds were weighed at the end of the food restriction period, and the length of this period was increased only if the bird’s weight remained stable (fluctuations less than ∼0.5g) for 4 days. Experiments began once birds had stable weight on 3-5 hours of food restriction and were observed caching into the replica cache sites in their home cages.

#### Caching task

Each bird was recorded in one 1-hour long behavioral session per day, starting immediately after the food restriction period. During these session, all four feeders in the arena were stocked with chopped sunflower seeds (∼25 mg fragments). At any moment in time, the arena could be in one of 5 states. In state 0, all feeders were closed. In states 1-4, the corresponding feeder (1-4) was open, while the other three feeders were closed. In most sessions, the state of the arena at the beginning of the session was chosen randomly with probability 1/4 of choosing state 0 and 3/16 of choosing any of the other states. A timer was also started, with duration chosen randomly from an exponential distribution between 60 and 90 s with 15 s decay. Once the timer duration elapsed, a new state was chosen, with the same probabilities: 1/4 for state 0 and 3/16 for the others. The timer was restarted with a new randomly chosen duration. A given state could repeat more than once, with one exception: if state 0 has occurred three times in a row, states 1-4 were chosen randomly with 1/4 probability each. In some early sessions, state 0 was omitted. In these sessions, the arena could be in states 1-4, and all transition probabilities between states were 1/4. For some birds, the timer duration was also chosen from smaller values: exponential distribution between 30 and 60 s with 15 s decay.

In most session, we also turned lights off periodically, which helped motivate food seeking and caching behaviors. In these cases, 5-min lights-on periods were alternated with lights-off periods, which lasted 0.5, 1, or 2 min in different sessions. In early sessions for each bird, the lights-off period was omitted.

Except for the analysis in Supplementary Figure 3A,C, we only analyzed data from those 10 birds that made at least 64 caches across all sessions. For the analysis in Supplementary Figure 3A,C, all 17 birds were used. For all analyses, we only included sessions where birds made at least 10 caches. For each bird, sessions were pooled across all feeder opening/closing and lights on/off conditions that were used for that bird.

#### Retrieval task

Each bird was recorded in one behavioral session per day, which lasted at least 30 min (see below). The session consisted of trials, each of which contained three phases: the feeder-open phase, the delay phase, and the feeder-closed phase.

The feeder-open phase consisted of two periods: the “ eating” period and the “ caching” period. In the eating period, one of the feeders was open, while the other three were closed. A timer was started at the beginning of this period, with random duration chosen from an exponential distribution between 30 and 60 s with 15 s decay. Once the timer had elapsed, a new feeder was chosen to be open with 1/4 probability, with the other three feeders closed; the same feeder could be repeated more than once. The timer was restarted with a new randomly chosen duration.

The caching period started whenever the bird cached the first seed – i.e., making one of the sites in the arena occupied. When this first cache was made, a new feeder was immediately chosen to be open with 1/4 probability. Subsequently, a new feeder was chosen with 1/4 probability whenever a new site was occupied by a cache, allowing the same feeder to be repeated more than once. This was done to motivate caching throughout the arena, rather than in the vicinity of the same feeder. When the bird visited the open feeder after caching the first seed, a 15 s timer was started. The feeder-open phase of the trial was terminated whenever this timer elapsed or 3 sites in the arena became occupied, whichever came first.

In the delay phase, all four feeders were closed, and lights in the arena were turned off. The delay phase lasted 2 min. After this 2 min period, lights were turned back on, and the feeder-closed phase was initiated.

In the feeder-closed phase, all four feeders remained closed at all times. If the bird retrieved all of the caches that it made earlier in the same trial, the feeder-closed phase terminated 15 sec after the removal of the last cache. This 15 sec window was implemented to accommodate possible recaching of the last retrieved seed. The feeder-closed phase also terminated after 10 min if the bird had not retrieved all of the caches that were made in the same trial. After the feeder-closed phase terminated, a new trial was immediately started if the duration of the session had not yet exceeded 30 min. The session was also terminated after 30 min if the bird never made any caches.

Data from all 7 birds were used in the analysis of this task.

#### Experiment controlling for olfactory cues

Birds were recorded in 3 sessions per day. During the first session (habituation session), all feeders were closed, and the bird had no access to food. The bird’s location in the arena was tracked for 5 min to measure the “ baseline” location distribution. Next, the bird was removed from the arena, then immediately reintroduced for another session (caching session). In the caching session, all feeders were open and the bird was allowed to make 3 caches. After 3 caches were made, the bird was again removed from the arena and placed into a cage with no food for a ∼30 min delay period. During this delay, the feeders were closed and all food was removed from the arena, including the bird’s caches and any scattered food. After the delay, the bird was reintroduced into the arena for the final session (test session). During this session, the bird’s location was again tracked for 5 min.

### Quantification and statistical analysis

#### Annotating site interactions

Using the bottom camera, we defined an ROI around each cache site. Pixel values from the first 100 frames of the video, when all cache sites were empty, were averaged to obtain a “ baseline” image for each ROI. For each subsequent frame in the video, we then measured two values: the Pearson correlation with the baseline image (*R*_b_), and the Pearson correlation with the previous frame (*R*_p_). *R*_b_ was normalized by subtracting the median and dividing by the standard deviation. *R*_p_ was transformed by subtracting a moving average of 200 frames. The MATLAB *findpeaks* function was used to detect changes in correlation that occurred in either one of these values (*R*_b_: minPeakDistance 100, minPeakHeight 10; *R*_p_: minPeakDistance 100, minPeakHeight 0.3). All of these events were visualized with side and bottom cameras and manually classified as caches, retrievals, checks, or false positives. Any period of time where the lights of the arena were turned off was excluded during this analysis.

#### Location tracking with deep neural networks

To determine the bird’s location, we trained a deep neural network^26,27^ to track two locations on the bird’s body using the top camera: the tip of the beak and the location halfway between the two feet. We then defined an ROI around each feeder, as well as around the perch adjacent to each cache site. A feeder visit was defined as the bird’s beak entering the corresponding ROI while the feeder was open. A cache site visit was defined as the bird’s feet entering the corresponding ROI.

#### Spatial distribution of caches and checks

Spatial distributions of caches (Caching task) and checks (Retrieval task) were quantified by computing the *p*^bias^ values (see Probabilistic model of behavior). To calculate whether these distributions differed from a uniform distribution, we measured the entropy of *p*^bias^ as

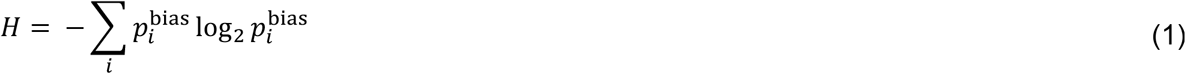

For our arena, *H* could range between 0 bits for a maximally biased bird that uses only using one site) to 6 bits for perfectly uniform behavior. To assess statistical significance, we compared this value of *H* to values from 10000 simulations, in which the same number of events was drawn randomly from a uniform distribution spanning all sites in the arena.

To determine whether *p*^bias^ was specific to individual birds, we divided each bird’s sessions into two groups (A and B) and assigned each session to one of these groups. The two groups were equal-sized if the number of sessions was even, or differed by one session if the number of sessions was odd. We computed distributions of cache or check locations separately for the two groups (*p*^bias,A^ and *p*^bias,B^), using the same formula as for *p*^bias^. We then computed the Pearson correlation between *p*^bias,A^ and *p*^bias,B^ for each bird and defined “ within-bird correlation” as the median of these values across birds. To compute “ across-bird correlations”, we shuffled the identities of all birds 1000 times and computed the median Pearson correlation between *p*^bias,A^ and *p*^bias,B^ across these 1000 shuffles. The entire process was repeated for 1000 shuffled datasets, in which the assignment into groups A and B was randomized.

For the “ within PC1 cluster” analyses in Figure 2D and Supplementary Figure 1D, we smoothed all values of *p*^bias^ with a 3×3 site Gaussian window (σ = 7.1cm, i.e. the distance between neighboring sites). We computed the first principal component of all smoothed *p*^bias^ values across birds. We grouped all birds into two clusters, based on whether the value of the first principal component was <0 or >0. We then performed the same process as above, but with one exception: during the shuffling of bird identities, a bird was only allowed the identity of another bird from the same cluster.

To measure the stability of *p*^bias^ time, we measured the distribution of cache or check locations for each trial separately, using the same formula as for *p*^bias^. We smoothed these distributions with a 3×3 site Gaussian window (σ = 7.1cm). We then measured the Pearson correlation value across all pairs of smoothed distributions that were Δ*s* sessions apart and took the median across all of these pairs of sessions. This value was computed for all Δ*s* from 1 to *N*. Here, *N* was determined as the value for which in the Caching task there were at least 2 pairs of sessions available from at least 5 of the birds. This value was *N* = 8, which was applied to both tasks.

### Probabilistic model of behavior

Consider a bird recorded in a set of behavioral sessions within the arena. This arena contains multiple discrete *sites*. Each site is either a *cache site* (i.e., location covered by a silicone flap) or a *feeder*. We will consider events that occur at these sites.

We first consider cache sites. At a cache site, a *check* is an opening of the silicone rubber flap by the bird, which allows the bird visual access to the contents of the site. A *cache* is a placement of seeds into the site. A *retrieval* is a removal of seeds from the site. Caches and retrievals necessarily require the bird to open the silicone flap; therefore, each cache and retrieval at a cache site is also considered to be a check. For feeders, we define a *retrieval* as a removal of a seed. A retrieval is the only type of an event that is considered at feeders.

We define a site *interaction* as any cache, check, or retrieval – either at a cache site or at a feeder. Visits to sites that do not include at least one of these three types of events (e.g., landing at a site) are not considered to be interactions and are not included in the analysis.

For each bird, the experiment consists of non-overlapping *trials*. In the Caching task, each behavioral session is considered to be one trial. In the Retrieval task, each trial is a period of time consisting of the *feeder-open* phase, the *delay* phase, and the *feeder-closed* phase. In this task, each behavioral session contains at least one trial, and usually multiple trials.

Each site is considered to be *occupied* if it contains at least one seed and *empty* if it does not. Feeders have a large capacity and are occupied at all times. Cache sites are considered to be empty at the beginning of each trial. If some cache sites contain seeds remaining from a previous trial, those seeds are ignored in the analysis. This occasionally happens in the Retrieval task if the bird had failed to retrieve all of the caches during one of the previous trials. Any withdrawal of such a seed is not considered to be a retrieval. However, any subsequent placement of that seed into a cache site is considered to be a cache.

#### Definitions of variables

*The following variables are defined for each bird:*

*S* is the number of sites in the arena. In the arena described in this paper, there are 64 cache sites and 4 feeders. Therefore, *S* = 68.

*R* is the number of behavioral trials.

*I* is the number of site interactions performed by the bird across all *R* trials.

*For each site interaction i, where* 1≤ *i* ≤ *I, the following are defined:*

*s*_*i*_ is the index of the site where the interaction occurred, where 1≤ *s*_*i*_ ≤ *S*.

*r*_*i*_ is the index of the trial during which the interaction occurred, where 1≤ *r*_*i*_ ≤ *R*.

*ϕ*_*i*_ is the phase of the trial during which the interaction occurred, defined as *ϕ*_*i*_ = 0 for all site interactions during the Caching task, *ϕ*_*i*_ = 1for interactions during the feeder-open phase of the Retrieval task, and *ϕ*_*i*_ = 2 for interactions during the feeder-closed phase of the Retrieval task. There are no interactions during the delay phase of the Retrieval task.

Δ_*i*_ is the change in the number of seeds at site *s*_*i*_ during the interaction. I.e., Δ_*i*_ < 0 for retrievals,

Δ_*i*_ > 0 for caches, and Δ_*i*_ = 0 for checks that are not coincident with either a cache or a retrieval.

*For each site s, where* 1≤ *s* ≤ *S, the following is defined:*

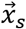 is the location of the site. Here, 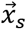 is a vector indicating location in 2-dimensional space. Several additional variables are defined for convenience of notation.

The Euclidean distance between sites *s* and *s*′ is:

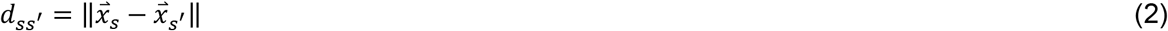

Next, we consider the time point immediately preceding interaction *i*. Several variables are defined to indicate the state of the behavioral arena at this point. For the following definitions, “ current trial” is the trial that contains interaction *i*.

*n*_*is*_ is the occupancy of site *s*. The value is *n*_*is*_ = 1if the site is occupied, or *n*_*is*_ = 0 if the site is empty.

N_*i*_ is the total occupancy of the arena – i.e., the number of cache sites that are occupied.

*t*_*is*_ is the amount of time that has elapsed since the most recent interaction with site *s* in the current trial. If the bird has not interacted with site *s* in the current trial, then *t*_*is*_ = ∞.

*c*_*is*_ indicates whether site *s* is “ checked-empty”. That is, *c*_*is*_ = 1if the bird has checked site *s* in the current trial, and that site is empty. If the bird has not checked site *s* in the current trial, or if site *s* is occupied, then *c*_*is*_ = 0.

*C*_*i*_ is the number of checked-empty cache sites in the arena.

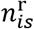 indicates whether the latest cache into site *s* in the current trial has been a recache. That is, 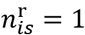 if the bird has cached into site *s* during the feeder-closed phase of the task, and site *s* is occupied. If the bird has only cached into site *s* during the feeder-open phase, or if site *s* is empty, then 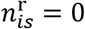.

*μ*_*i*_ indicates whether the next interaction in the current trial that involves a seed is a cache. If the next interaction involving a seed is a cache, *μ*_*i*_ = 1. If the next such interaction is a retrieval, or if the bird never interacts with a seed again in the current trial, *μ*_*i*_ = 0.

#### Model parameters

Twelve parameters are used by various models described in the main text. We define a vector of these parameters:

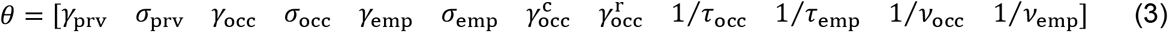

Inversions of the last four parameters are done for the purpose of parameter regularization, described later.

#### Interaction masks

Models described in the text are applied to specific subsets of interactions. A single model is sometimes applied to different subsets: for example, Model #1 is used to quantify the effect of proximity either on caches or on checks, which are two different subsets of interactions. To define such subsets, we use *interaction masks*. For each interaction *i*, the value of the mask is 1 if that interaction is included in the analysis or 0 if it is not. For different formulas below, two masks will be used: *G*_*i*_ is used to mask interactions based on the task (Caching or Retrieval); *H*_*i*_ is used to further mask subsets of interactions within these tasks.

**Table 1.** Masks for different subsets of interactions.

| Subset of interactions | $G_i$ | $H_i$ |
| --- | --- | --- |
| All caches in the Caching task | $G_i = \begin{cases} 1, & \phi_i = 0, \Delta_i > 0 \\ 0, & \text{otherwise} \end{cases}$ | $H_i = \begin{cases} 1, & \phi_i = 0, \Delta_i > 0 \\ 0, & \text{otherwise} \end{cases}$ |
| First checks in the Caching task. This mask includes the first check that the bird made after retrieving each seed that was subsequently cached (i.e., it does not include checks made after retrieving seeds that were subsequently eaten without caching). | $G_i = \begin{cases} 1, & \phi_i = 0, \Delta_i > 0 \\ 0, & \text{otherwise} \end{cases}$ | $H_i = \begin{cases} 1, & \phi_i = 0, \Delta_{i-1} < 0, \mu_i = 1 \\ 0, & \text{otherwise} \end{cases}$ |
| All checks in the feeder-closed phase of the Retrieval task | $G_i = \begin{cases} 1, & \phi_i = 2 \\ 0, & \text{otherwise} \end{cases}$ | $H_i = \begin{cases} 1, & \phi_i = 2 \\ 0, & \text{otherwise} \end{cases}$ |
| All checks up to and including finding a cache (i.e., making the first check of an occupied site) in the feeder-closed phase of the Retrieval task. If no cache is found in the feeder-closed phase of the trial, this includes all checks up until the end of the trial. | $G_i = \begin{cases} 1, & \phi_i = 2 \\ 0, & \text{otherwise} \end{cases}$ | $H_i = \begin{cases} 1, & \phi_i = 2, \sum_{\substack{j < i \\ r_j = r_i}} n_{js_j} = 0 \\ 0, & \text{otherwise} \end{cases}$ |
| First checks in the feeder-closed phase of the Retrieval task. This mask includes the first check made during the feeder-closed phase of each trial. | $G_i = \begin{cases} 1, & \phi_i = 2 \\ 0, & \text{otherwise} \end{cases}$ | $H_{ik} = \begin{cases} 1, & \phi_{i-1} = 1, \phi_i = 2 \\ 0, & \text{otherwise} \end{cases}$ |

#### Scaling factors

In the baseline model (Model #0), the probability of an interaction occurring at a particular site given by the distribution *p*^bias^. We compute *p*^bias^ for either the Caching task or the Retrieval task using the mask *G* described above. For every site *s*:

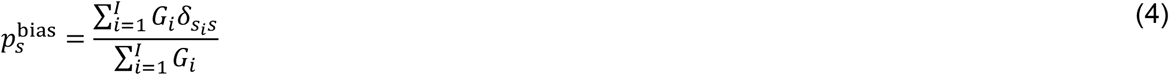

where *δ* is the Kronecker Delta, defined as *δ*_*xy*_ = 1if *x* = *y*, and *δ*_*xy*_ = 0 if *x* ≠ *y*.

We next define *scaling factors*. A scaling factor is a function defined for each interaction *i* and site *s*. These factors will be used later to scale values of *p*^bias^. Positive values of these scaling factors indicate an increase in probability, and negative values indicate a decrease in probability.

The first factor is used to model the effect proximity to the previous site interaction. It is a Gaussian that decays with distance from the previous site, defined at distances >0. Empirically, we found that models fit better if the value of this scaling factor was set to 0 at a distance of 0. This is due to the fact that birds were unlikely to interact with the exact same site twice in a row, even though they were highly likely to interact with neighboring site. Therefore, we defined the scaling factor as follows:

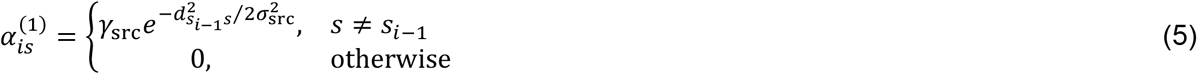

The second factor is used to model the effect of distance from occupied sites in the arena. It is a summation of Gaussians, each centered at an occupied site. The amplitude decays in time (in order to model memory decay) and with the occupancy of the arena (in order to model memory capacity). Note that in most models in the paper these decays are not included (i.e. *τ*_occ_ = ∞ and/or *v*_occ_ = ∞).

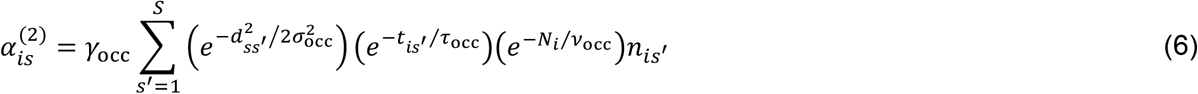

The third factor is analogous to the second factor, but for checked-empty sites in the arena:

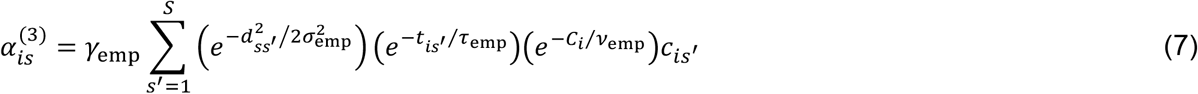

The fourth and fifth factors are used to separately model the effects of caches made in the feeder-open phase of the trial and caches made in the feeder-closed phase (i.e., “ recaches”):

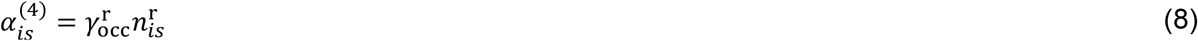

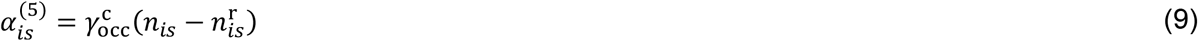

Each of the scaling factors *k* = 1,2, …, 5 is converted to a multiplicative form as follows:

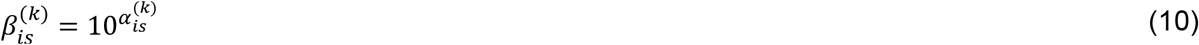

Un-normalized probability of interaction *i* occurring at site *s* is then computed by multiplying all five scaling factors by the baseline probability. Because this value depends on all model parameters, we denote it as a function of *θ*:

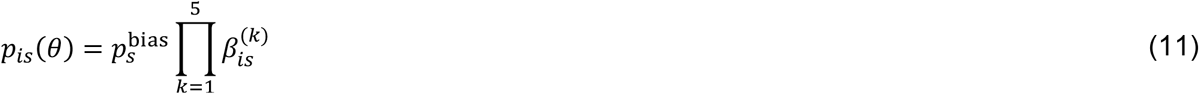

This value is converted to normalized probability that sums to 1 across all sites of the arena:

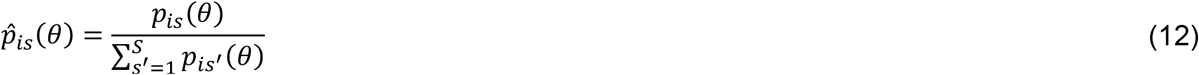

#### Model definitions

We describe twelve different models in the main text. All of these models can be expressed using the mathematical formulation described above. The difference between these models is the parameter space: In some models, a particular parameter is fixed at a value of 0, whereas in other models, that same parameter is a free parameter permitted to take values within a certain range. The following table specifies the parameter space for each model. Models are numbered using index *M* = 0,1, …, 11. Parameters within vector *θ* are numbered using index *z* = 1,2, …, 12. For each model *M* and parameter *z*, the table indicates *P*(*M, z*) – the set of permitted values for that parameter. For parameters whose values are fixed at zero, *P*(*M, z*) = {0}. We use *f*_*M*_ to indicate the number of free parameters in model *M*; this value is also given in the table.

**Table 2.**
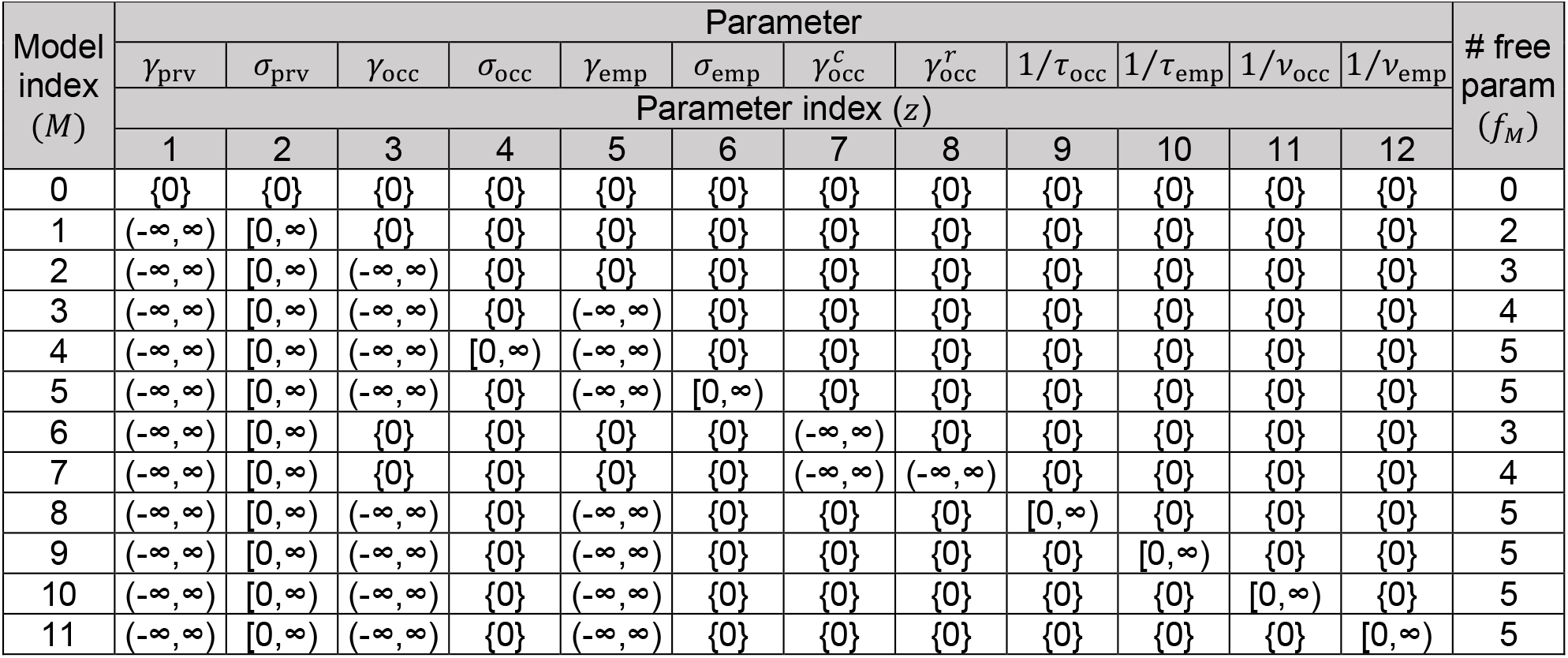
Permitted ranges of parameters, *P*(*M, z*), for different models.

| Model<br>index<br>($M$) | Parameter | | | | | | | | | | | | # free<br>param<br>($f_M$) |
| --- | --- | --- | --- | --- | --- | --- | --- | --- | --- | --- | --- | --- | --- |
| | $\gamma_{\text{prv}}$ | $\sigma_{\text{prv}}$ | $\gamma_{\text{occ}}$ | $\sigma_{\text{occ}}$ | $\gamma_{\text{emp}}$ | $\sigma_{\text{emp}}$ | $\gamma_{\text{occ}}^c$ | $\gamma_{\text{occ}}^r$ | $1/\tau_{\text{occ}}$ | $1/\tau_{\text{emp}}$ | $1/\nu_{\text{occ}}$ | $1/\nu_{\text{emp}}$ | |
| | Parameter index ($z$) | | | | | | | | | | | | |
|  | 1 | 2 | 3 | 4 | 5 | 6 | 7 | 8 | 9 | 10 | 11 | 12 |  |
| 0 | {0} | {0} | {0} | {0} | {0} | {0} | {0} | {0} | {0} | {0} | {0} | {0} | 0 |
| 1 | $(-\infty, \infty)$ | $[0, \infty)$ | {0} | {0} | {0} | {0} | {0} | {0} | {0} | {0} | {0} | {0} | 2 |
| 2 | $(-\infty, \infty)$ | $[0, \infty)$ | $(-\infty, \infty)$ | {0} | {0} | {0} | {0} | {0} | {0} | {0} | {0} | {0} | 3 |
| 3 | $(-\infty, \infty)$ | $[0, \infty)$ | $(-\infty, \infty)$ | {0} | $(-\infty, \infty)$ | {0} | {0} | {0} | {0} | {0} | {0} | {0} | 4 |
| 4 | $(-\infty, \infty)$ | $[0, \infty)$ | $(-\infty, \infty)$ | $[0, \infty)$ | $(-\infty, \infty)$ | {0} | {0} | {0} | {0} | {0} | {0} | {0} | 5 |
| 5 | $(-\infty, \infty)$ | $[0, \infty)$ | $(-\infty, \infty)$ | {0} | $(-\infty, \infty)$ | $[0, \infty)$ | {0} | {0} | {0} | {0} | {0} | {0} | 5 |
| 6 | $(-\infty, \infty)$ | $[0, \infty)$ | {0} | {0} | {0} | {0} | $(-\infty, \infty)$ | {0} | {0} | {0} | {0} | {0} | 3 |
| 7 | $(-\infty, \infty)$ | $[0, \infty)$ | {0} | {0} | {0} | {0} | $(-\infty, \infty)$ | $(-\infty, \infty)$ | {0} | {0} | {0} | {0} | 4 |
| 8 | $(-\infty, \infty)$ | $[0, \infty)$ | $(-\infty, \infty)$ | {0} | $(-\infty, \infty)$ | {0} | {0} | {0} | $[0, \infty)$ | {0} | {0} | {0} | 5 |
| 9 | $(-\infty, \infty)$ | $[0, \infty)$ | $(-\infty, \infty)$ | {0} | $(-\infty, \infty)$ | {0} | {0} | {0} | {0} | $[0, \infty)$ | {0} | {0} | 5 |
| 10 | $(-\infty, \infty)$ | $[0, \infty)$ | $(-\infty, \infty)$ | {0} | $(-\infty, \infty)$ | {0} | {0} | {0} | {0} | {0} | $[0, \infty)$ | {0} | 5 |
| 11 | $(-\infty, \infty)$ | $[0, \infty)$ | $(-\infty, \infty)$ | {0} | $(-\infty, \infty)$ | {0} | {0} | {0} | {0} | {0} | {0} | $[0, \infty)$ | 5 |

For each model, the parameter space is defined as the Cartesian product of permitted ranges across all twelve parameters:

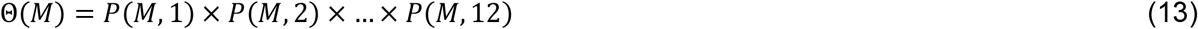

#### Log-likelihood computation

The log-likelihood of the model is computed by adding log-probabilities across all interactions observed in the data. For this purpose, the interaction mask *H* is used to only include the subset of the interactions that is being fit by the model:

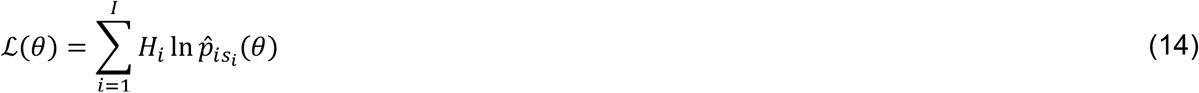

To pull data across birds, we denote the log-likelihood for each bird *b* as ℒ_*b*_(*θ*). Here 1≤ *b* ≤ *B*, where *B* is the number of birds. The pooled log-likelihood is then computed as:

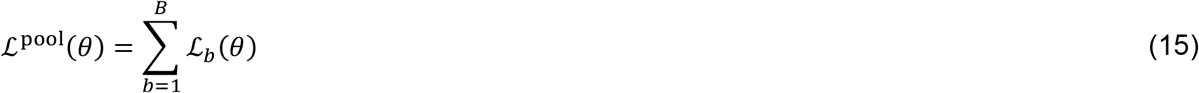

The “ cost” of the model fit is computed by negating the likelihood and adding a Ridge regularization term. Ridge regularization penalizes large magnitudes of parameters and helps prevent overfitting. For bird *b*, the cost is:

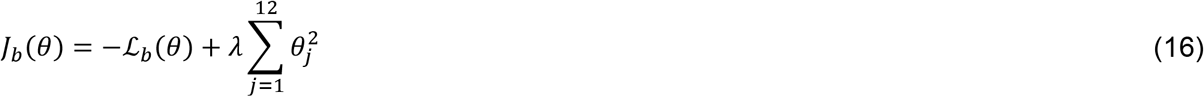

For the pooled data, the cost is similarly:

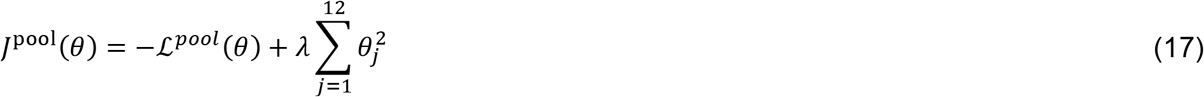

Here, *λ* is the regularization parameter. We use *λ* = 1 for the Caching task and *λ* = 10 for the Retrieval task. These values are different due to different numbers of data points in the two tasks.

#### Model fitting and evaluation

To fit each model *M* using maximum-likelihood estimation, we determine values within the parameter space of model *M* that minimize the cost. These best-fit parameters are given by 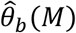 for each bird *b* and by 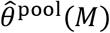 for the pooled data:

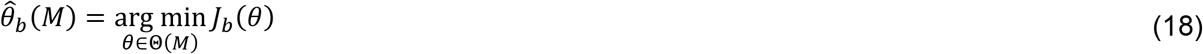

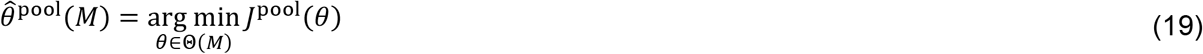

We fit each model using the MATLAB *fmincon* function. Each model was fit 5 times with different initial conditions, and the solution with the minimum cost was selected.

To evaluate each model, we used the Akaike Information Criterion (AIC). Smaller AIC values indicate better model performance. AIC value is improved by increased model likelihood, but is penalized by adding extra parameters. We computed the AIC value on the pooled data:

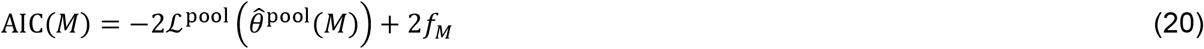

## Supplemental figures

**Figure S1.**
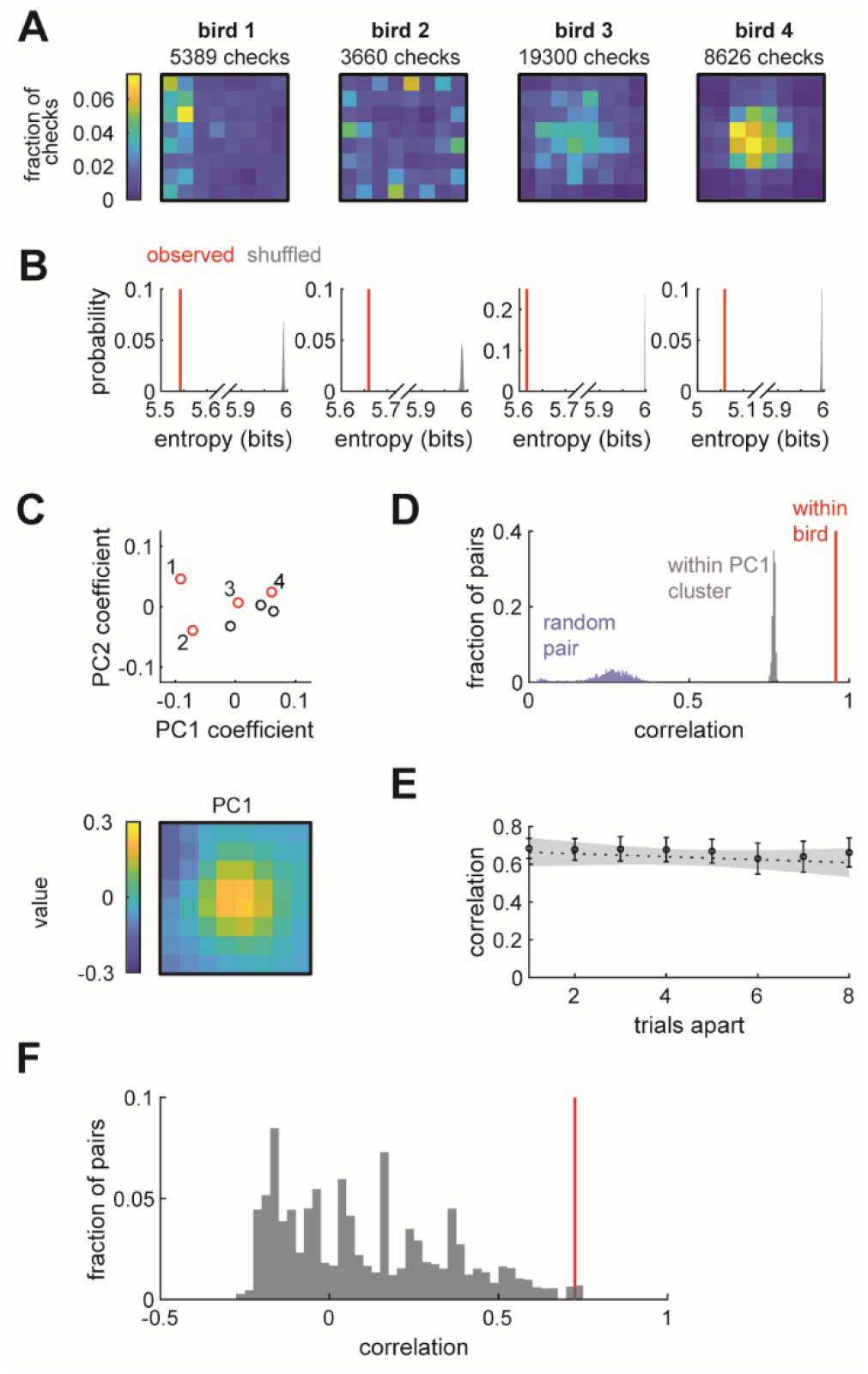
Chickadees exhibit idiosyncratic and stable biases in the locations of checks. Related to Figure 2. (A) Distributions of check locations across all sessions for four example birds. Distributions are denoted by p^bias^ in the text. (B) Red lines: entropy values of the spatial distributions shown in (A). Grey histograms: entropy values for simulated checks drawn from a uniform distribution. Number of simulated checks was the same as in the observed data. (C) Principal component analysis of p^bias^ values. Top: coefficients of the first two principal components for all birds. Red circles and numbers indicate birds shown in (A) and (B). Bottom: the first principal component. (D) Pearson correlation of p^bias^ between subsets of all sessions paired within bird, between different birds, and between birds selected from the same PC1 cluster shown in (C). Although birds do not visibly cluster in (C), we defined “ clusters” as groups of birds whose first principal component was <0 and >0 for analogy with Fig 2. (E) Pearson correlation of p^bias^ between pairs of sessions at different time lags. Values are medians across birds. Error bars: sem. Dashed line: linear regression. Grey shade: 95% confidence of linear regression. Slope of the regression was not significantly different from zero (p=0.39). (F) Pearson correlation of p^bias^ between all sessions of the Caching and all sessions of the Retrieval tasks. Red line: median within bird correlation. Grey histogram: correlation of randomly paired birds.

**Figure S2.**
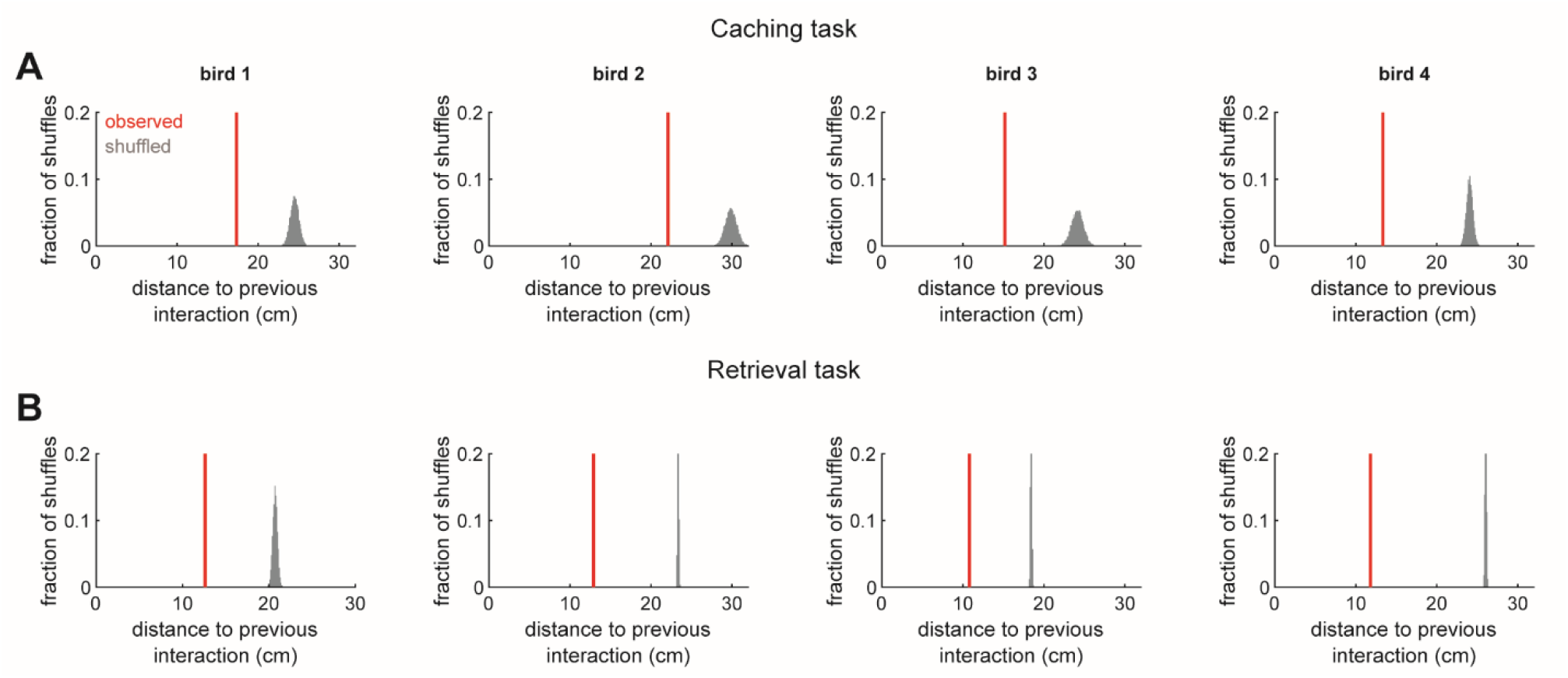
Corroboration of the proximity effect. Related to Figure 4. (A) Distance between the location of the check preceding a cache and the site of the cache for four example birds. Red: mean of observed distances. Grey: distribution of mean distances for shuffled distances, in which caches and checks were paired randomly. (B) Same as (A) but for the Retrieval task.

**Figure S3.**
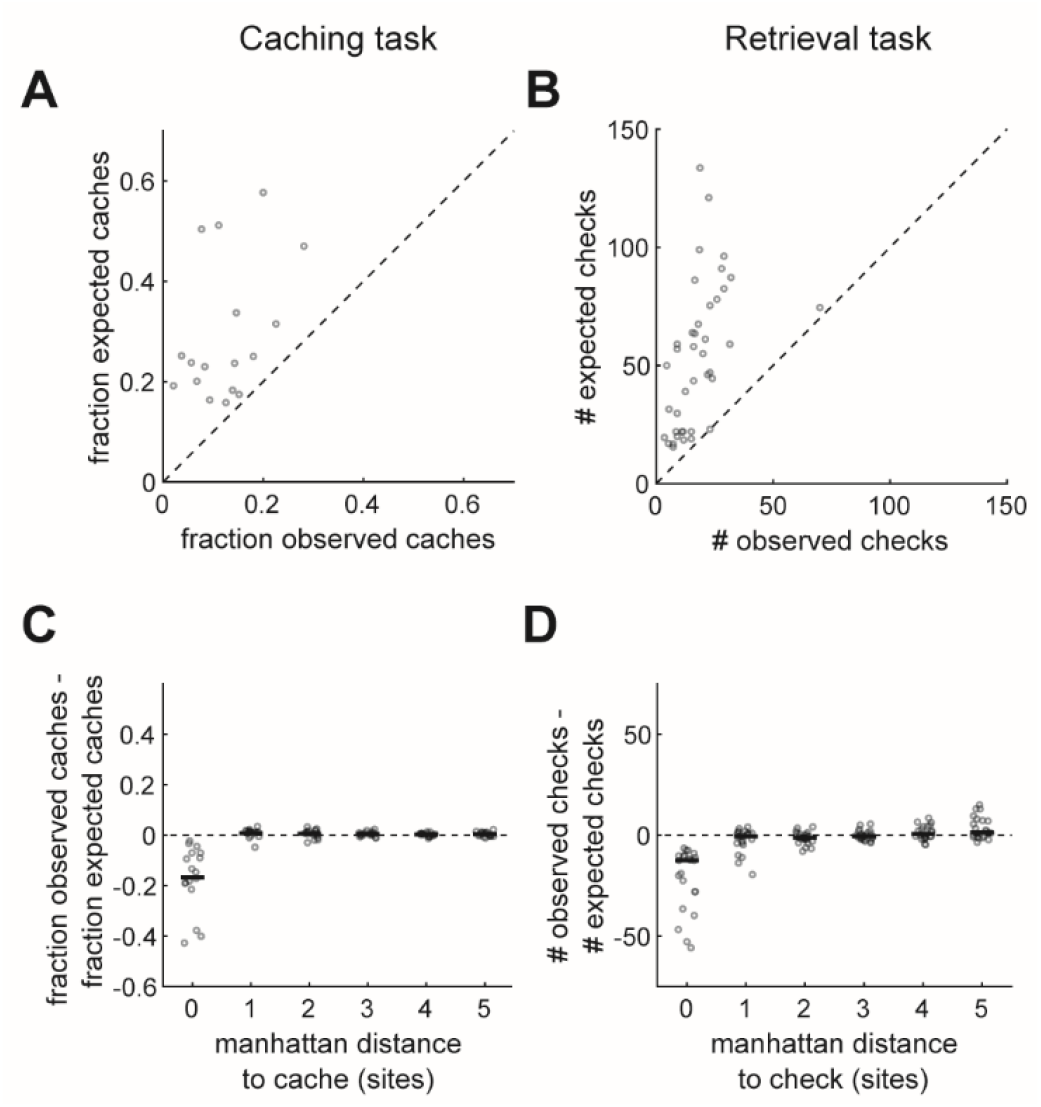
Corroboration of the effects of specific site contents on behavior. Related to Figure 5. (A) Observed and expected fractions of caches made into occupied sites. Each symbol indicates fractions for one bird. Expected fraction is calculated by randomly drawing, at the time of each cache, a simulated cache location from the distribution p^bias^. Random drawing was performed 1000 times, and fractions of occupied sites were averaged across the draws. (B) The number of observed and expected checks required by the bird to find a cache in the Retrieval task. For each cache, we ordered all trials of the same bird across all sessions and contatenated the sequences of checks made in those trials. For calculating the observed number of checks, the actual trial containing the cache was first in the order, and the remaining trials were ordered randomly. For calculating the expected number of checks, all trials were ordered randomly. In both cases, we asked where in the concatenated sequence the earliest check of the cache site occurred. The median across 1000 random orders was taken. For most caches, this analysis compared how many attempts the bird required to find a cache vs. how many attempts a trajectory taken from a different trial would require to find that same cache. However, in a small number cases when the bird failed to find the cache, this analysis equally penalized both the observed and the expected numbers. Each symbol is the median value across all caches made into a particular site (1-64), including only those caches sites where at least 2 birds cached at least 3 times each (40 sites total). (C) Difference between the observed and expected fractions of caches into occupied sites, calculated as in (A). For distance=0, this analysis uses occupied sites, as in (A). For distances>0, this analysis instead takes the average across all sites in the arena at the corresponding manhattan distance away from an occupied site. Black line indicates the median across all birds. Birds avoided caching into occupied sites, but did not avoid caching into sites even at a distance of 1 site away (i.e., the effect was site-specific). (D) Analysis described in (B), performed for different manhattan distances away from the cache site. For each site *j* in the arena, we considered the set of all sites *S*_*dj*_ that are located at a manhattan distance *d* from *j*. We then computed the median difference between observed and expected values for each of the sites in S_*dj*_ across all caches made into *j*. Finally, we took this median value from the site in *S*_*dj*_ that has the smallest absolute value. These values are plotted for the same 40 sites as in (B). Black line is the median across all sites. Birds showed attraction to occupied sites in the Retrieval task, but did not show attraction even at a distance of 1 site away (i.e., the effect was also site-specific).

**Figure S4.**
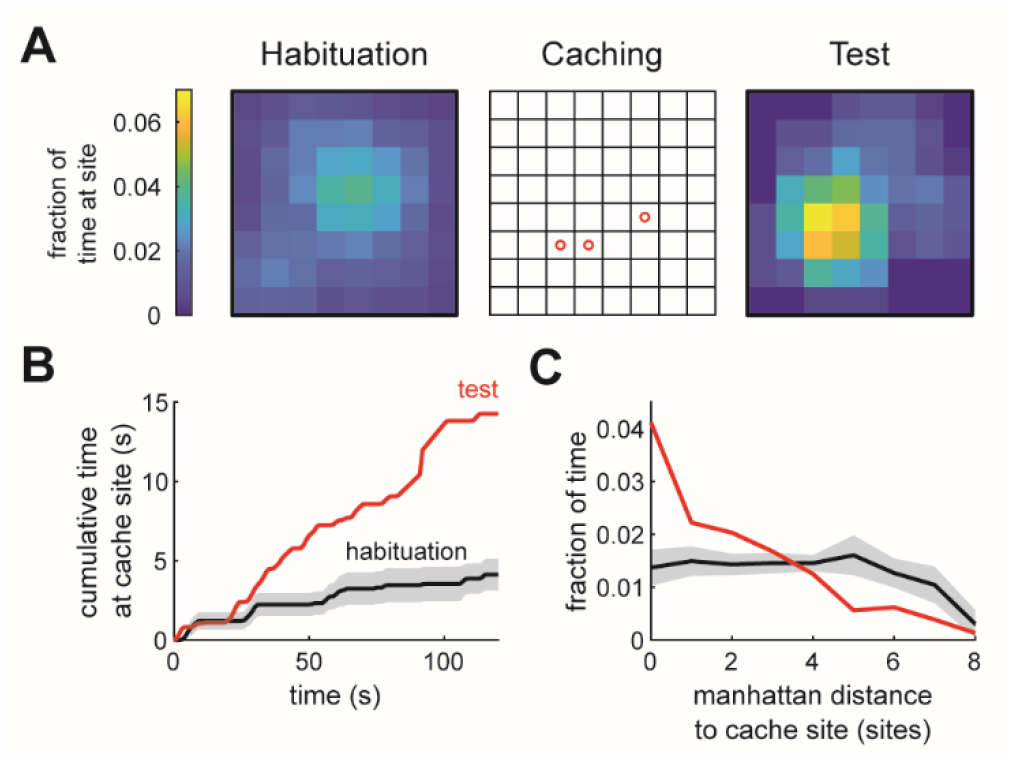
Chickadees return to cache locations without seeds present. Related to Figure 6. (A) Example behavioral session in the experiment controlling for olfactory cues. Left: occupancy of the chickadee in the first 5 min of the habituation session. Center: locations of caches made in the caching session. Right: spatial occupancy in the first 60 s of the test session. For the left and right panel, any uninterrupted time spent >10 s at one site (such as during eating or grooming) was excluded. (B) Cumulative time spent at the cache site before caching (habituation session) and after caching (test session). Red trace: mean across all test sessions (n = 6) in all birds. Black trace: mean across all habituation sessions. Grey: ± sem across sessions. Time 0 is the beginning of the session. (C) Fraction of time spent at each cache site in first 120 s of the session, as a function of manhattan distance away from the nearest cache site. Red trace: mean across all sessions in all birds. Grey: ±sem across sessions. Effect is observed not only at the cache site, but also at nearby sites because the bird often spent time at a neighboring site visually inspecting the site where a cache was expected.

